# α_1_ adrenergic receptor - PKC - Pyk2 - Src signaling boosts L-type Ca^2+^ channel Ca_v_1.2 activity and long-term potentiation in rodents

**DOI:** 10.1101/2022.07.01.498400

**Authors:** Kwun Nok Mimi Man, Peter B. Henderson, Karam Kim, Mei Shi, Mingxu Zhang, Madeline Nieves-Cintron, Manuel F. Navedo, Mary C. Horne, Johannes W. Hell

**Affiliations:** Department of Pharmacology, University of California, Davis, CA 95616-8636, USA.; Department of Pharmacology, University of Iowa, Iowa City, IA 52242-1109, USA.

## Abstract

The cellular mechanisms mediating norepinephrine functions in brain to result in behaviors are unknown. We identified the L-type Ca^2+^ channel (LTCC) Ca_V_1.2 as a principal target for G_q_- coupled α1-adrenergic receptors (ARs). α_1_AR signaling increased LTCC activity in hippocampal neurons. This regulation required PKC-mediated activation of the tyrosine kinases Pyk2 and, downstream, Src. Pyk2 and Src were associated with Ca_V_1.2. In model neuroendocrine PC12 cells, stimulation of PKC induced tyrosine phosphorylation of Ca_V_1.2, a modification abrogated by inhibition of Pyk2 and Src. Upregulation of LTCC activity by α_1_AR and formation of a signaling complex with PKC, Pyk2, and Src suggests that Ca_V_1.2 is a central conduit for signaling by norepinephrine. Indeed, a form of hippocampal LTP in young mice requires both the LTCC and α_1_AR stimulation. Inhibition of Pyk2 and Src blocked this LTP, indicating that enhancement of Ca_V_1.2 activity via α_1_AR - Pyk2 - Src signaling regulates synaptic strength.

## INTRODUCTION

Norepinephrine (NE) causes arousal and augments behavioral acuity and learning (*1–5*). NE signals via the G_q_ - coupled α_1_ adrenergic receptor (AR), G_i_ - coupled α_2_AR, and G_s_ – coupled β_1_ and β_2_ARs. β_1_ and β_2_ARs act through adenylyl cyclase (AC), cAMP, and PKA (*6*). The β_2_AR, G_s_, AC, and PKA are all associated with the L-type Ca^2+^ channel (LTCC) Ca_v_1.2 for efficient signaling in neurons (*7–12*) and heart (*13*). The formation of this signaling complex identifies Ca_v_1.2 as a major effector of signaling by NE. We now find that Ca_v_1.2 is also a major effector for signaling via the α_1_AR, which has a higher affinity for NE than βARs (*14, 15*). Importantly a large body of evidence implicates the α_1_AR in NE’s role in attention and vigilance (*16–22*).

Ca_v_1.2 fulfills a remarkably broad spectrum of functions. Dysfunctions due to mutations in Ca_v_1.2 span from impaired cardiac contractility to the autistic-like behaviors seen in Timothy syndrome (*23*). Furthermore, Ca_v_1.2 has been linked to filopodia formation in invasive cancer cells (*24*). Ca_v_1.2 is by far the most abundant LTCC in heart and accounts for ∼80% of all LTCCs in brain (*25, 26*). It governs the heartbeat, vascular tone and neuronal functions including long-term potentiation (*11, 12, 27-29*), long-term depression (*30*), neuronal excitability (*31, 32*), and gene expression (*9, 33–38*). Studies on Ca_V_1.2 mutant mice suggest that this channel plays a central role in anxiety disorders, depression, and self-injurious behavior (*26*). Congruently, LTCC blockers elicit antidepressant effects while agonists induce depression-like behavior (*39, 40*) and self-biting in mice, a symptom associated with autism (*41*).

Ca_V_1.2 consists of the pore-forming subunit α_1_1.2, a β subunit and the α_2_δ subunit (*42–44*). The β and α_2_δ subunits facilitate release of α_1_1.2 subunits from the endoplasmic reticulum, inhibit ubiquitin-mediated degradation of voltage-gated calcium channels, influence electrophysiological properties of Ca^2+^ channels, such as activation and inactivation, and play diverse roles in the regulation of these channels (*42–44*).

In the cardiovascular system the α_1_AR, the endothelin receptor ET1, and the angiotensin receptor AT_1_ are important regulators of LTCC currents via G_q_ signaling (*42, 45*). G_q_ stimulates phospholipase Cγ to induce production of diacylglycerol (DAG) and inositol-1,4,5-trisphosphate (IP_3_), which triggers Ca^2+^ release from intracellular stores. DAG and Ca^2+^ act in concert with phosphatidyl-serine (PS) to activate different PKC isoforms. Stimulation of PKC mostly leads to an increase in Ca_V_1.2 activity (*43, 45–50*). However, an inhibitory effect of PKC on Ca_V_1.2 currents has been reported in cardiomyocytes (*51*). This inhibition is mediated by phosphorylation of residues T27 and T31 by PKC in an isoform of α_1_1.2 that is expressed in heart (*52*). T27/T31 are not present in the most prevalent brain isoform due to alternative splicing (*53*), thus the inhibitory effect of PKC on LTCC currents is typically absent in neurons and neural crest-derived PC12 cells, or in vascular smooth muscle (*50, 54*). Here we show that stimulation of the α_1_AR and of PKC consistently augments LTCC in hippocampal neurons.

Despite the prominent role of PKC in augmentation of Ca_V_1.2 activity, how PKC mediates this effect has been unknown. PKC activates the nonreceptor tyrosine kinase Pyk2, a signaling process first shown in PC12 cells (*55, 56*) and later primary neurons (*57, 58*), and cardiomyocytes (*59*). Activation of PKC triggers autophosphorylation of residue Y402 on Pyk2 to create a binding site for the SH2 domain of Src, which upon binding to Pyk2 becomes activated (*55*). Src increases LTCC activity in smooth muscle cells (*60–62*), retinal pigment epithelium (*63*), and neurons (*62, 64, 65*). Furthermore, PKC (*66, 67*) and Src (*60, 65, 68*), are physically and functionally associated with Ca_V_1.2. These findings underscore the physiological relevance of Src in regulating Ca_V_1.2. Importantly, the pathway by which Src is activated in the context of Ca_V_1.2 regulation has not been determined.

Once we established that stimulation of PKC or the G_q_/PKC – coupled α_1_AR strongly augments LTCC activity in neurons, we tested whether Pyk2 mediates this upregulation of channel activity. We link the α_1_AR – PKC signaling to Src, and define it as the terminal tyrosine kinase phosphorylating Ca_V_1.2 downstream of G_q_-coupled receptors. In neurons, the nearly 2-fold increase in LTCC currents upon stimulation of PKC with PMA or via the α_1_AR was blocked by inhibitors of Pyk2 and Src, consistent with earlier data showing that Src elevates Ca_V_1.2 activity to a comparable degree (*62, 65*). Furthermore, we found that Pyk2 coimmunoprecipitated with Ca_V_1.2 in parallel to Src. We identified the loop between domains two and three of α_1_1.2 as the Pyk2 binding site. Stimulation of PKC either directly with phorbol-12-myristate-13-acetate (PMA) or through the G_q_-coupled bradykinin receptor leads to tyrosine phosphorylation of α_1_1.2 in PC12 cells. Abrogation of Pyk2 or Src activity ablated the phosphorylation. Finally, we discovered that the long-term potentiation (LTP) in young mice mediated by LTCC-dependent Ca^2+^ influx during 200 Hz tetani (termed LTP_LTCC_) that is not NMDAR-dependent, required α_1_AR stimulation and both Pyk2 and Src activity. These findings implicate upregulation of Ca_v_1.2 activity by α_1_AR - Pyk2 - Src signaling as a critical process for control of synaptic strength. Our findings indicate that Ca_V_1.2 forms a supramolecular signaling complex (signalosome) with PKC, Pyk2, and Src and that α_1_AR – PKC – Src – Ca_V_1.2 signaling constitutes a central regulatory mechanism of neuronal activity and synaptic plasticity by NE.

## RESULTS

### α1AR signaling augments LTCC activity in hippocampal neurons via PKC, Pyk2, and Src

We performed cell-attached recordings from cultured hippocampal neurons for single-channel analysis, which allows pharmacological isolation of LTCCs by application of ω-conotoxins GVIA and MVIIC (*10–12, 69*). Application of phenylephrine (PHE), a selective agonist for all three α_1_ARs, augmented open probability of LTCCs from 0.18±0.0433 (H_2_O vehicle Control, n=11) to 0.6156±0.1386 (PHE; n=13, p≤0.01; Fig. 1A,B). This increase was blocked by the selective α_1_AR antagonist prazosin (0.2954±0.0607; n=10, p≤0.05), indicating that PHE acted through α_1_ARs and not other G protein-coupled receptors. Prazosin by itself had no effect, versus vehicle control (0.1846±0.04624; n=10) suggesting that there is little if any regulation of LTCCs under basal conditions in neurons by α_1_ARs. PHE also increased the peak current of the ensemble average current in a prazosin-sensitive manner (Fig. 1C,D). Our cell-attached recordings report NPo, i.e., the product of channel number N times open probability Po.

**Figure 1.**
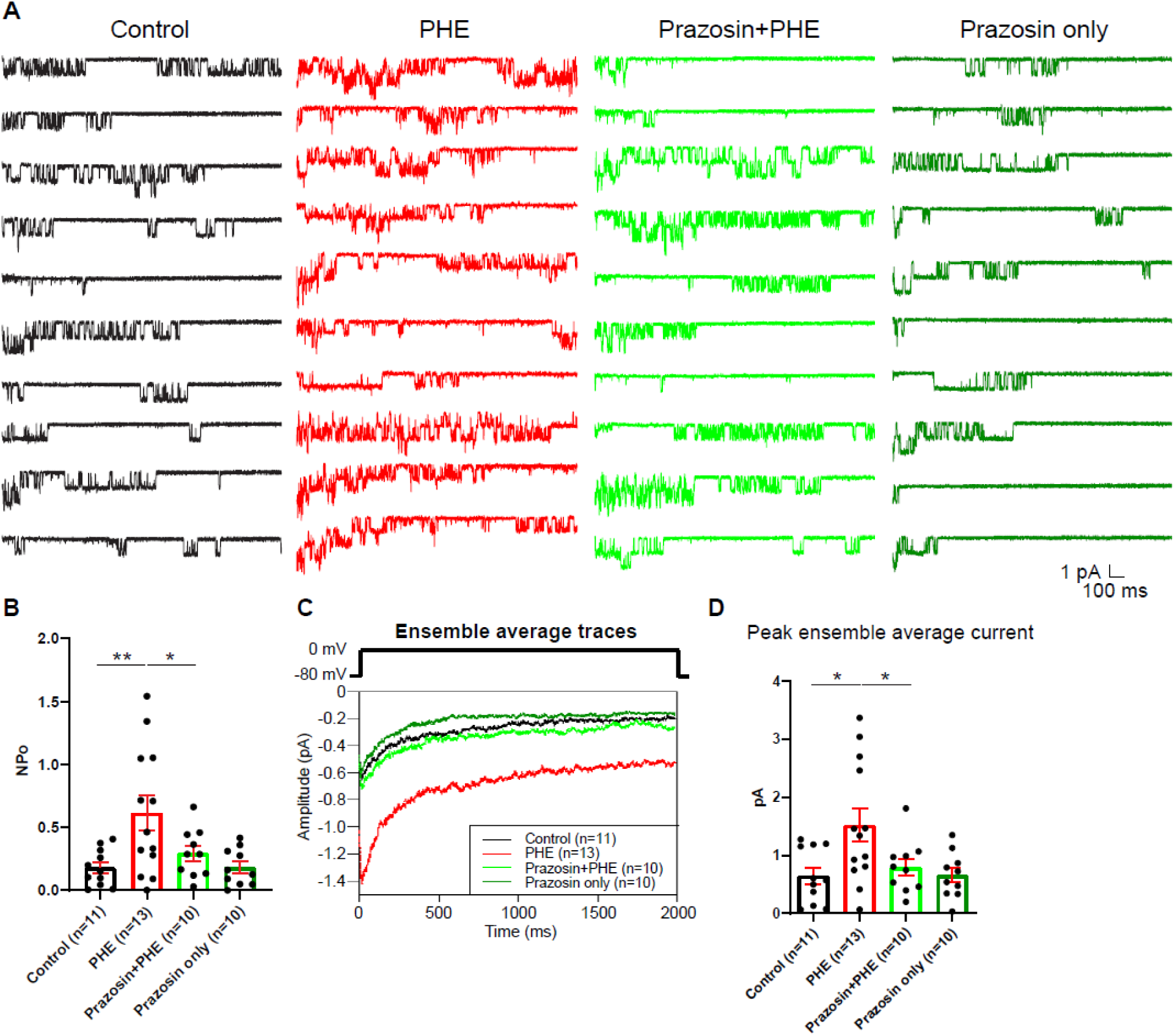
The α_1_AR agonist phenylephrine (PHE) augments NPo of L-type Ca^2+^ channels (LTCC) in hippocampal neurons. **(A)** Ten consecutive traces from representative cell-attached single-channel recordings of LTCCs from cultured hippocampal neurons with vehicle (water; black), 10 µM PHE (red), PHE plus 20 nM prazosin (bright green), and prazosin alone (dark green). **(B)** The increase in NPo by PHE was blocked by prazosin. F_3,40_=5.474. Control vs. PHE, *P*=0.0036; PHE vs. Prazosin+PHE, *P*=0.0334; Control vs. Prazosin only, *P*=0.9723. **(C)** Ensemble averages during depolarization. **(D)** The increase in ensemble average peak currents by PHE was blocked by prazosin. F_3,40_=4.506. Control vs. PHE, *P*=0.0101; PHE vs. Prazosin+PHE, *P*=0.0316; Control vs. Prazosin only, *P*=0.9722. **(B,D)** Data are presented as means ± SEM. n represents the number of cells (\**P*≤0.05, \*\**P*≤0.01; ANOVA with post-hoc Holm-Sidak’s multiple comparisons test).

However, in several experiments PHE was applied after seal formation, which resulted in the same increase in NPo (Man and Hell, unpublished). Thus, we interpret the PHE-induced increase of NPo as an increase in channel open probability Po because it appears impossible that the number of channels under the patch pipette would change during that experiment.

Because direct phosphorylation of α_1_1.2 by PKC inhibits Ca_V_1.2 activity (*52*), we explored whether PKC might upregulate Ca_V_1.2 activity indirectly via other kinases. PKC can activate Pyk2 (*55–58*) and thereby Src (*55, 57*). Src, in turn, augments LTCC activity (*60–65*). Therefore, we tested whether block of Pyk2 and Src affects upregulation of LTCC activity by PHE. In a new set of recordings augmentation of LTCC activity by PHE from NPo of 0.2008±0.03348 (DMSO vehicle control, n=33) to 0.3272±0.04412 (PHE, n=33; p≤0.01; Fig. 2A,B) was completely blocked by two different PKC inhibitors, bisindolylmaleimide I (GF109203X; Bis I; 0.1412±0.03305; n=11, p≤0.01) and chelerythrine (Chel; 0.05801±0.01508; n=12, p≤0.001), the Pyk2-selective inhibitor PF-719 (0.09118±0.02828; n=10, p≤0.01) and two structurally different Src family kinase inhibitors, PP2 (0.01487±0.006808; n=7, p≤0.001) and SU6656 (0.09149±0.02866; n=9, p≤0.01). Peak currents of ensemble averages showed respective changes (Fig. 2C,D). Accordingly, α_1_AR signaling increases LTCC activity via a PKC-Pyk2-Src signaling cascade. Notably, stimulation of other major G_q_ protein - coupled receptors in neurons, i.e., the metabotropic mGluR1/5 receptors with dihydroxyphenylglycine (DHPG) and muscarinic receptors with muscarine, did not significantly increase LTCC activity (Fig. S1).

**Figure 2.**
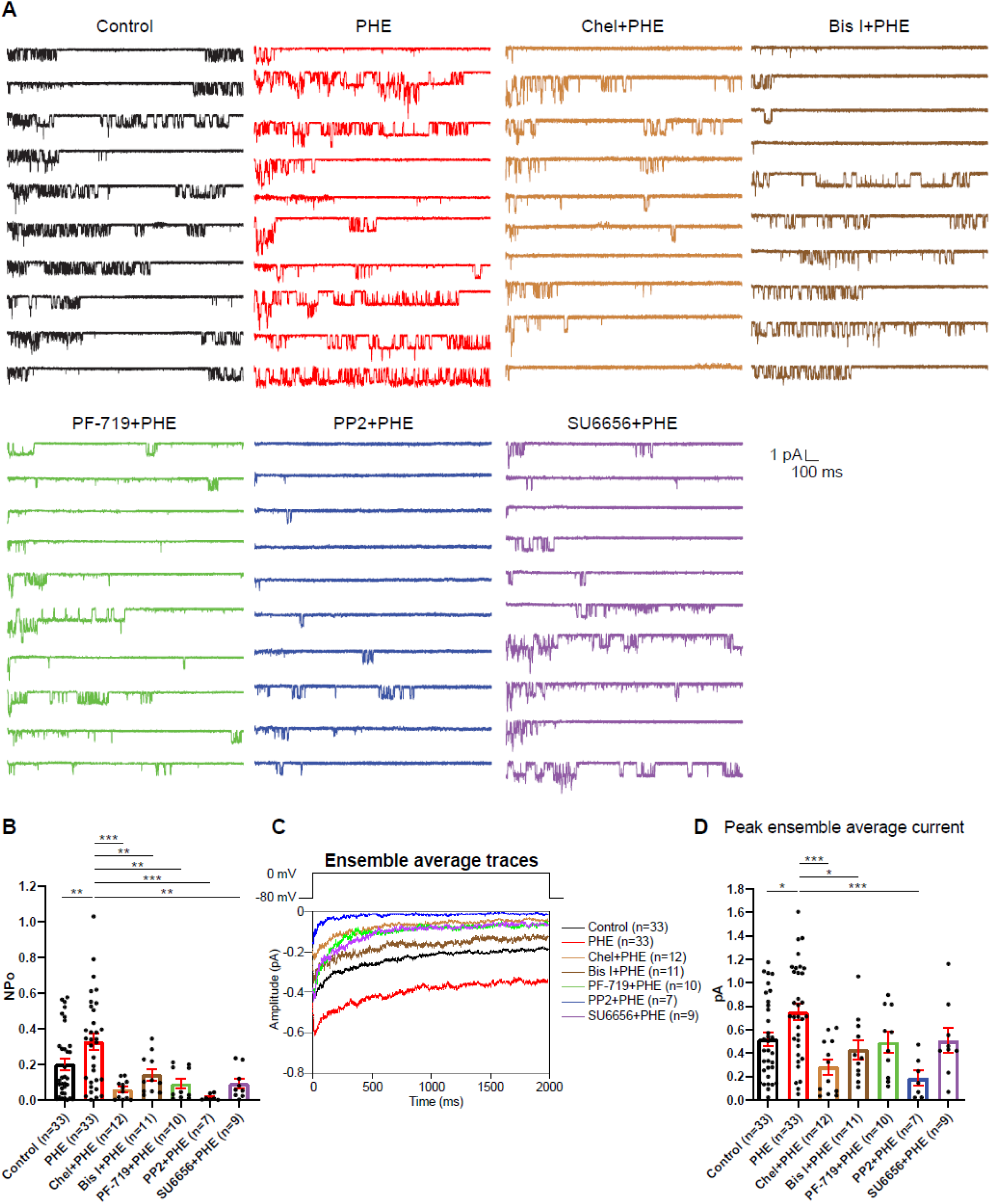
The PHE-induced increase in NPo of LTCCs in hippocampal neurons requires PKC, Pyk2, and Src. **(A)** Ten consecutive traces from representative cell-attached single-channel recordings of LTCCs with vehicle (0.1% DMSO; black) and PHE either alone (red) or with the PKC inhibitors chelerythrine (Chel; 10 µM; bright brown) and bisindolylmaleimide I (Bis I; 100 nM; dark brown), the Pyk2 inhibitor PF-719 (1 µM; green), or the Src inhibitors PP2 (10 µM; blue) and SU6656 (10 µM; purple). **(B)** The increase in NPo by PHE was blocked by all inhibitors. F_6,108_=6.434. Control vs. PHE, *P*=0.0076; PHE vs. Chel+PHE, *P*=0.0001; PHE vs. Bis I+PHE, *P*=0.0076; PHE vs. PF- 719+PHE, *P*=0.0018; PHE vs. PP2+PHE, *P*=0.0003; PHE vs. SU6656+PHE, *P*=0.0022. **(C)** Ensemble averages during depolarization. **(D)** The increase in ensemble average peak currents by PHE was blocked by PKC inhibitors chelerythrine, bisinolylmaleimide I and Src inhibitor PP2. F_6,108_=4.839. Control vs. PHE, *P*=0.0242; PHE vs. Chel+PHE, *P*=0.0004; PHE vs. Bis I+PHE, *P*=0.0242; PHE vs. PF- 719+PHE, *P*=0.0723; PHE vs. PP2+PHE, *P*=0.0006; PHE vs. SU6656+PHE, *P*=0.0723. **(B,D)** Data are presented as means ± SEM. n represents the number of cells (\**P*≤0.05, \*\**P*≤0.01, \*\*\**P*≤0.001; ANOVA with post-hoc Holm-Sidak’s multiple comparisons test).

### PKC augments LTCC activity in hippocampal neurons via Pyk2 and Src

To further establish a role of Pyk2 and Src, we directly stimulated PKC by including PMA in the bath solution during single-channel recording of LTCCs in hippocampal neurons. PMA increases NPo of LTCCs by around two-fold from 0.1099±0.0173 (DMSO control, n=45) to 0.232±0.03269 (PMA, n=38; p≤0.001, Fig. 3A,B). This increase is blocked by Pyk2 inhibitors PF-719 (NPo=0.1407±0.02705, n=13, p≤0.05) and PF-431396 (NPo=0.09282±0.01765, n=18, p≤0.001) and by Src inhibitors PP2 (NPo=0.05614±0.01815, n=14, p≤0.001) and SU6656 (NPo=0.02951±0.00555, n=8, p≤0.001). Peak currents of ensemble averages showed respective changes (Fig. 3C,D). The L-type calcium channel blocker isradipine completely blocked L-type currents in the presence of PMA, indicating successful isolation of L-type single-channel currents (Fig. S2). These results show that in hippocampal neurons, PKC activation stimulates LTCC activity and this augmentation requires Pyk2 and Src activity.

**Figure 3.**
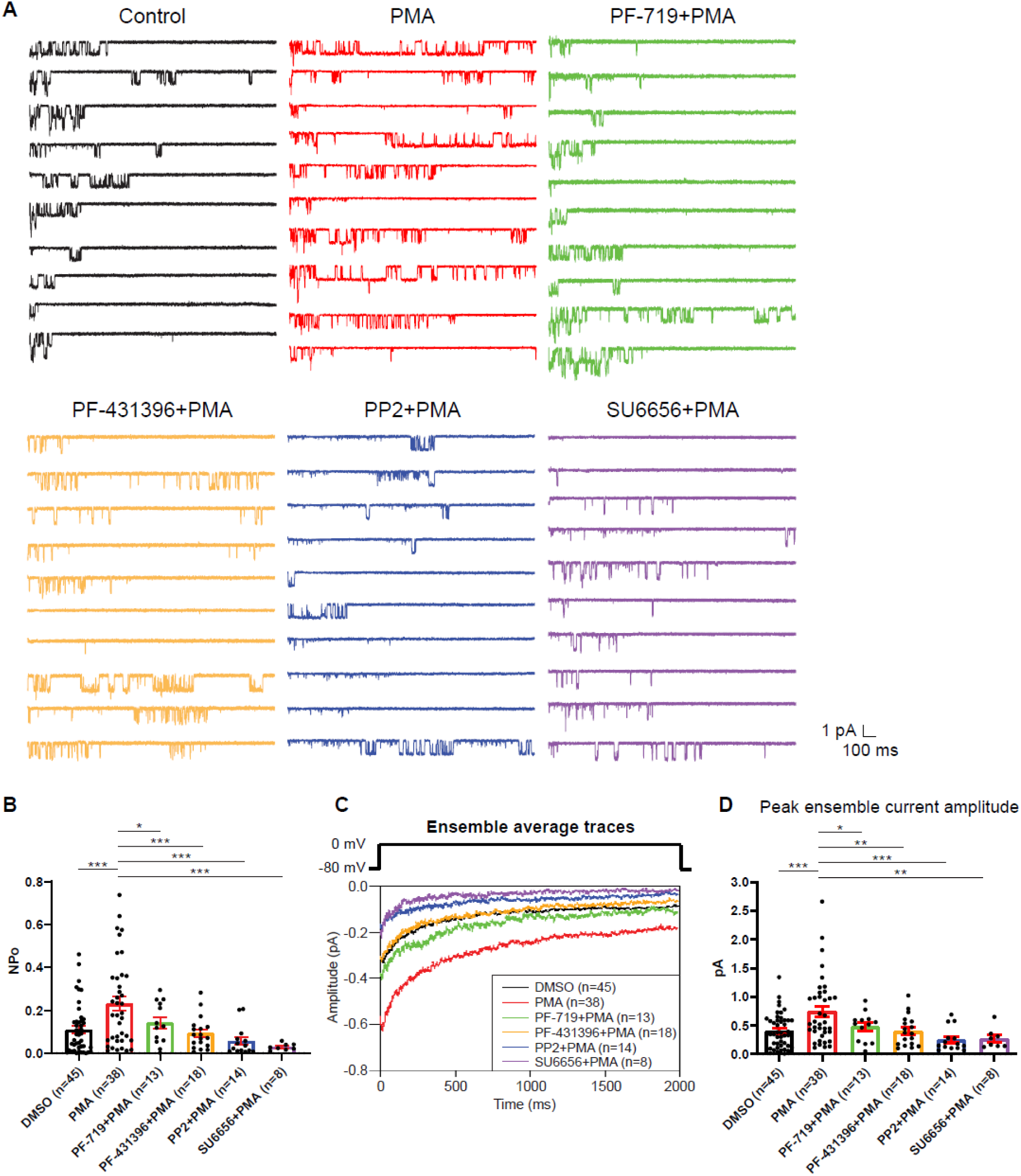
The increase in NPo of LTCCs in hippocampal neurons by PKC requires Pyk2 and Src. **(A)** Ten consecutive traces from representative cell-attached single-channel recordings of LTCCs with vehicle (0.06% DMSO; black) and 2 µM PMA either alone (red) or with the Pyk2 inhibitors PF-719 (1 µM; green) and PF-431396 (3 µM; orange), or the Src inhibitors PP2 (10 µM; blue) and SU6656 (10 µM; purple). **(B)** The increase in NPo by PMA was blocked by all inhibitors. F_5,130_=6.530. DMSO vs. PMA, *P*=0.0003; PMA vs. PF-719+PMA, *P*=0.0372, PMA vs. PF-431396+PMA, *P*=0.0009; PMA vs. PP2+PMA, *P*=0.0003; PMA vs. SU6656+PMA, *P*=0.0005. **(C)** Ensemble averages during depolarization. **(D)** The increase in ensemble average peak currents by PMA was blocked by all inhibitors. F_5,130_=5.665. DMSO vs. PMA, *P*=0.0003; PMA vs. PF-719+PMA, *P*=0.0303, PMA vs. PF- 431396+PMA, *P*=0.0051; PMA vs. PP2+PMA, *P*=0.0003; PMA vs. SU6656+PMA, *P*=0.0051. **(B,D)** Data are presented as means ± SEM. n represents the number of cells (\**P*≤0.05, \*\**P≤*0.01, \*\*\**P*≤0.001; ANOVA with post-hoc Holm-Sidak’s multiple comparisons test).

### Pyk2 co-immunoprecipitates with Ca_V_1.2 from brain and heart

Kinases and their regulators are often found in complexes with their ultimate target proteins (i.e., their signaling relay partners and ultimate substrates) including different ion channels for efficient and specific signaling (*43, 70*). Both, PKC (*66, 67*) and Src (*60, 65, 68*), are associated with Ca_V_1.2. We tested in brain and heart (where Ca_V_1.2 is most abundant) whether the same is true for Pyk2. The Pyk2 antibody detected a single immunoreactive band with an apparent M_R_ of

∼120 kDa in brain lysate (Fig. 4A) and two bands in the same range in heart (Fig. 4A,B), as reported earlier (*71*). The shorter form is missing 42 residues in the proline-rich region of Pyk2, which affects its binding selectivity to proteins with SH3 domains. The single size form of Pyk2 present in brain and its two size forms expressed in heart co-immunoprecipitated with Ca_V_1.2 (Fig. 4A,B). No Pyk2 immunoreactive band was detectable when the immunoprecipitation (IP) was performed with control IgG, demonstrating that the co-IP of Pyk2 with Ca_V_1.2 was specific. The detergent extracts from brain and heart were cleared of non-soluble material by ultracentrifugation prior to co-IP of Pyk2 with Ca_V_1.2. Thus, our findings indicate that Pyk2 forms a bona fide protein complex with Ca_V_1.2 rather than just co-residing in a detergent-resistant subcellular compartment.

**Figure 4.**
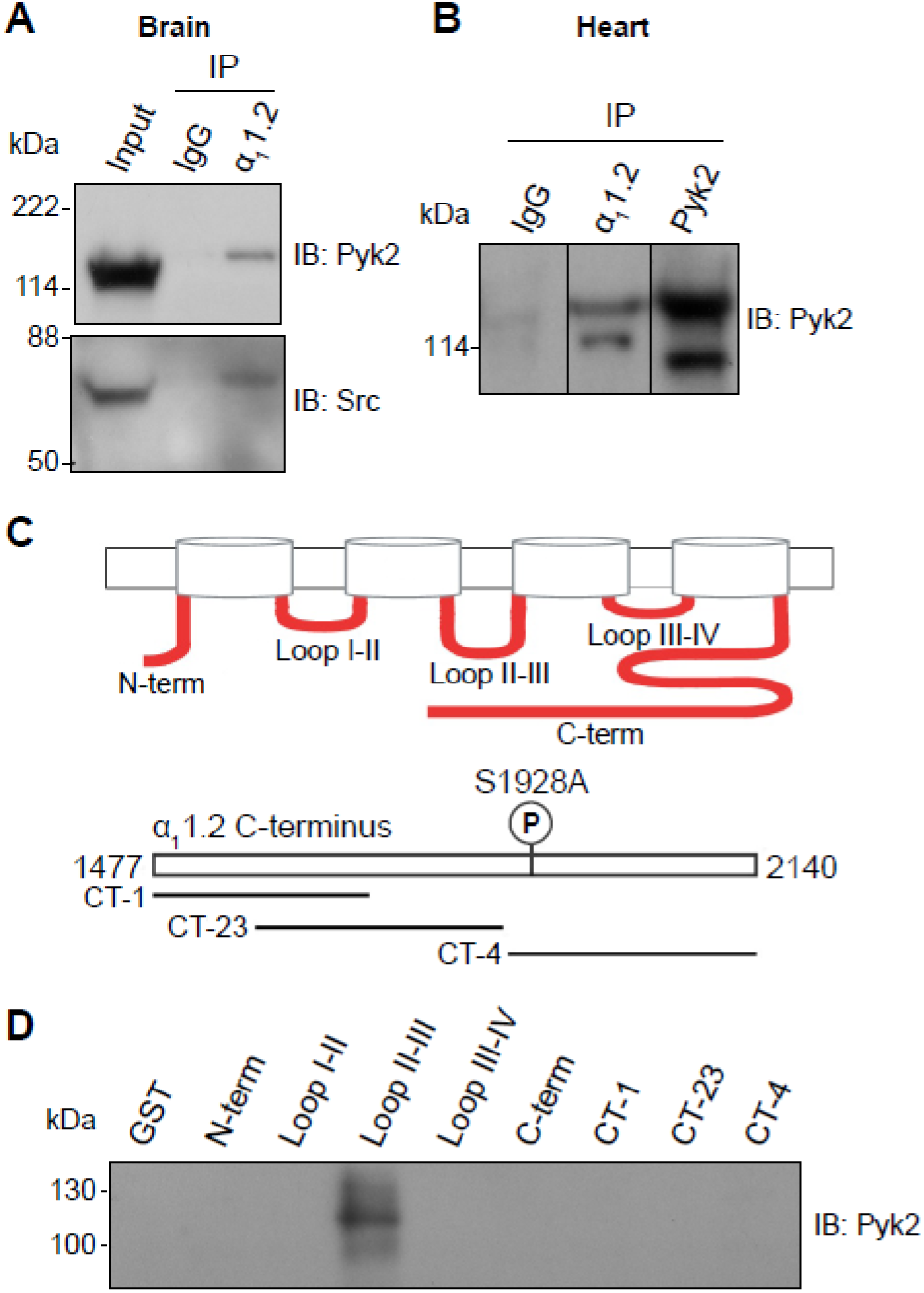
Pyk2 binds to the loop between domains II and III of α_1_1.2. **(A,B)** Co-immunoprecipitation of Pyk2 and Src with Ca_V_1.2 from brain (A) and heart (B). Triton X-100 extracts were cleared from non-soluble material by ultracentrifugation before immunoprecipitation (IP) with antibodies against α_1_1.2, Pyk2 itself, or non-immune control antibodies (IgG) and immunoblotting (IB) with anti-Pyk2 and anti-Src. Brain lysate (A, Input; 20 μl) and Pyk2 immunoprecipitates (B) served as positive control for detection of Pyk2 and Src by IB. Lanes for IgG control and α_1_1.2 IP in B are from the same IB and exposure but rearranged to eliminate non-relevant lanes. Pyk2 IP in B is from the same IB but is shown as a shorter exposure than the IgG and α_1_1.2 IP lanes. Comparable results were obtained in 4 independent experiments. **(C)** Schematic diagram of the intracellular α_1_1.2 fragments used in the pulldown assay (Table S1). **(D)** Pulldown assay of Pyk2 binding to α_1_1.2 fragments. GST fusion proteins of the N-terminus, the loops between domains I and II, II and III, III and IV, the whole C-terminus, and three different overlapping fragments covering the C-terminus were expressed in *E. coli*, immobilized on glutathione Sepharose, washed and incubated with purified His-tagged Pyk2. Comparable amounts of fusion proteins were present (data not shown but see (*11, 69, 97, 139*)). Comparable results were obtained in 5 independent experiments.

### Pyk2 binds to the loop between domains II and III of α_1_1.2

To further confirm a direct interaction between Pyk2 and Ca_V_1.2 we performed pull-down experiments using purified solubilized His-tagged Pyk2 and purified bead-bound GST-tagged α_1_1.2 fragments covering all intracellular regions of α_1_1.2 (Table S1). As demonstrated earlier all α_1_1.2 fragments were present in comparable amounts (*11, 69*). The GST fusion protein covering the loop between domains II and III of α_1_1.2 specifically pulled down Pyk2 (Fig. 4C,D) indicating that Pyk2 directly binds to this region of the α_1_ subunit.

### Inhibitors of Pyk2 and Src block the increase in α_1_1.2 tyrosine phosphorylation upon stimulation of PKC

PC12 cells are of neural-endocrine crest origin and widely used as model cells for neuronal signaling and development. They express high levels of Ca_V_1.2 (*54, 72–74*) and Pyk2 (*55, 56, 58*), making them an ideal model system for the difficult biochemical analysis of Ca_V_1.2 phosphorylation. To characterize tyrosine phosphorylation of α_1_1.2 we performed IP with the general anti-phosphotyrosine antibody 4G10 (*75, 76*). For this purpose, lysates were extracted with 1% SDS at 65°C followed by neutralization of SDS and ultracentrifugation before IP with the general anti-phosphotyrosine antibody 4G10 (*75, 76*). IP with 4G10 followed by immunoblotting (IB) with antibodies against the protein of interest is more reliable and more broadly applicable than the inverse. Because α_1_1.2 does not re-associate with its binding partners after complex dissociation with SDS and the neutralization and dilution of SDS with Triton X-100 (*77–79*) (see also (*80*)), detection of α_1_1.2 by IB in the 4G10 IP would reflect specific tyrosine phosphorylation of the α_1_1.2 subunit and not its artefactual re-association with an α_1_1.2 – associating tyrosine phosphorylated protein that had been pulled down by the 4G10 antibody. This approach also allows analysis of tyrosine phosphorylation of Pyk2 within the same sample.

PC12 cells were pretreated with vehicle (0.02% DMSO), the Pyk2 inhibitor PF-431396, or the Src inhibitors SU6656 and PP2 or its inactive analogue PP3 for 5 min before application of bradykinin or PMA for 10 min. Bradykinin strongly activates Pyk2 in PC12 cells via its G_q_- coupled cognate receptor (*55, 56*). Using the 4G10 IP method we found that both PMA and bradykinin increased tyrosine phosphorylation of Pyk2 as previously (*55, 56*) (Fig. 5A,B). This increase was prevented by PF-431396. In parallel, we determined phosphorylation of Pyk2 on Y402 and Y579 by direct IB of PC12 lysates with corresponding phosphospecific antibodies. Upon stimulation via PKC, Pyk2 phosphorylates itself in trans on Y402 (*58, 81*) and then binds with phosphoY402 to the SH2 domain of Src (*55*). This binding stimulates Src (*55*), which in turn phosphorylates Pyk2 on Y579 in its activation loop for full activation (*82*). Src also phosphorylates itself in trans on Y416 in its activation loop for its own full activation (*83*), which was determined in parallel with a phosphospecific antibody. We found that bradykinin and PMA increased phosphorylation of Pyk2 on Y402 (Fig. 5A,C) and Y579 (Fig. 5D,E) and of Src on Y416 (Fig. 5F,G). PF-431396 blocked all of these phosphorylations indicating that Pyk2 acts downstream of PKC and upstream of Src. Furthermore, the Src inhibitor PP2, but not its inactive analogue, PP3, also prevented PMA-induced Src autophosphorylation on Y416 (Fig. 5H,I), as expected. Finally, PP2 inhibited PMA-induced phosphorylation of Pyk2 on Y402 and Y579 indicative of a self-maintaining positive feedback loop between Pyk2 and Src (Fig. 5J-L). These results support the specific activation of both, Pyk2 and Src under our conditions and suggest that this activation occurs in a self-sustaining manner, which creates a quasi-molecular memory.

**Figure 5.**
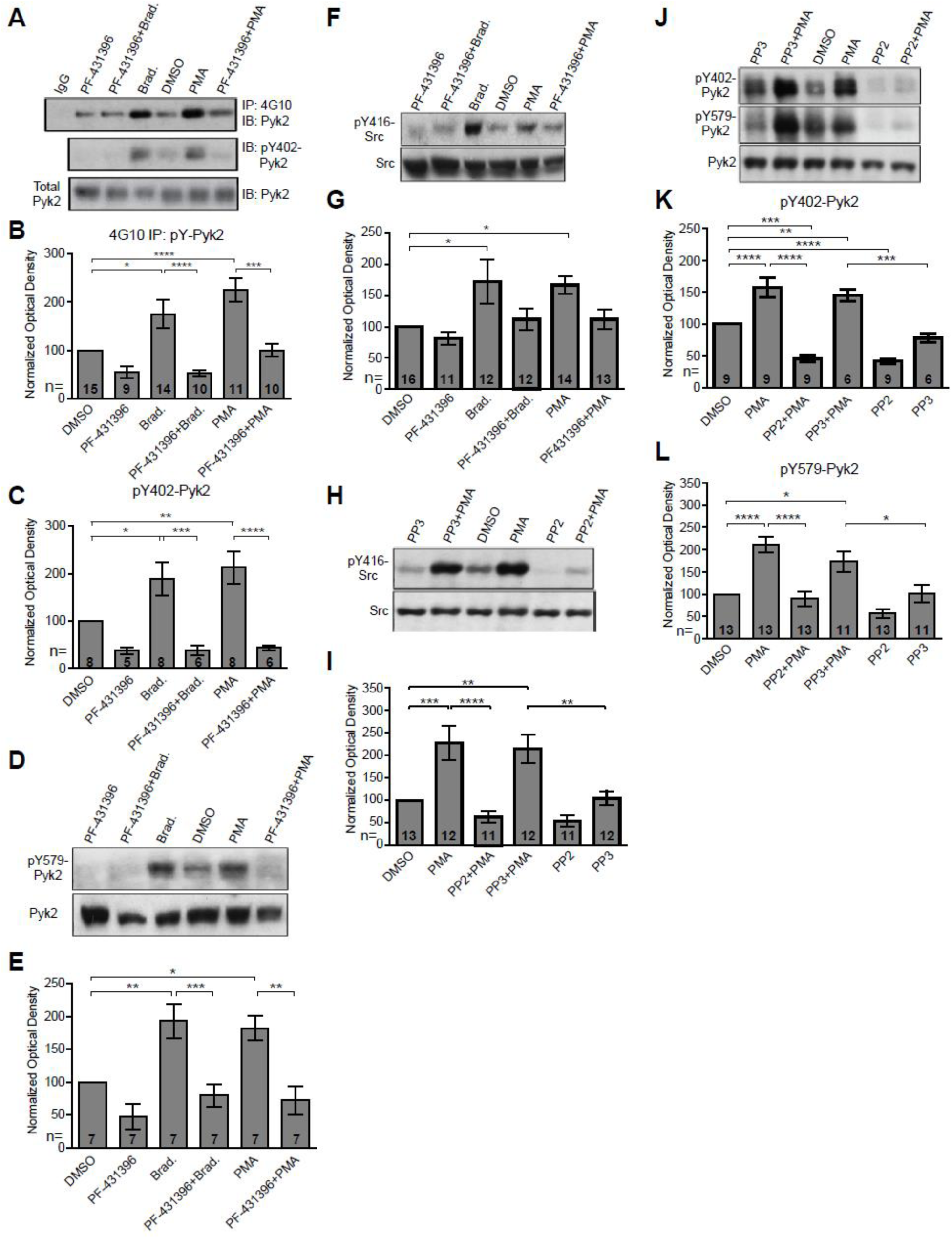
PKC activates interdependent Pyk2 and Src. PC12 cells were pretreated with vehicle (0.02% DMSO), the Pyk2 inhibitor PF-431396 (3 μM), and the Src inhibitor PP2 (10 μM) or its inactive analogue PP3 (10 μM) for 5 min before application of bradykinin (Brad., 2 μM) or PMA (2 μM) for 10 min, extraction with 1% SDS at 65°C to ensure dissociation of all proteins, neutralization of SDS with excess of Triton X-100, and ultracentrifugation. Supernatants were analysed by direct IB with the indicated Pyk2 and Src antibodies. Some samples underwent IP with the anti-phosphotyrosine antibody 4G10 before IB with anti-Pyk2 antibody (top panel in A and quantification in B). IgG indicates control IP with non-immune mouse IgG. **(A)** Upper panel: total pY levels of Pyk2 determined by IP with 4G10 and IB with anti-Pyk2. Middle panel: pY402 levels of Pyk2 detected with anti-pY402 in corresponding lysates. Lower panel: Levels of total Pyk2 detected with anti-Pyk2 in same lysates. **(B)** Ratios of total pY of Pyk2 after 4G10 IP to total Pyk2 in lysates, normalized to control. F_5,63_=12.73. DMSO vs. Brad., *P*=0.012; DMSO vs. PMA, *P*<0.0001; Brad. vs PF-431396+Brad., *P*<0.0001; PMA vs. PF-431396+PMA, *P*=0.0001. **(C)** Ratios of pY402 to total Pyk2 signals in lysates, normalized to control. F_5,35_=10.94. DMSO vs. Brad., *P*=0.039; DMSO vs. PMA, *P*=0.0052; Brad. vs PF-431396+Brad., *P*=0.0005; PMA vs. PF-431396+PMA, *P*<0.0001. **(D)** Upper panel: pY579 levels of Pyk2 detected with anti-pY579. Lower panel: Levels of total Pyk2 detected with anti-Pyk2 in same lysates. **(E)** Ratios of pY579 to total Pyk2 signals in lysates, normalized to control. F_5,36_=10.18. DMSO vs. Brad., *P*=0.0072; DMSO vs. PMA, *P*=0.021; Brad. vs PF-431396+Brad., *P*=0.0008; PMA vs. PF-431396+PMA, *P*=0.0011. **(F,H)** Upper panels: pY416 levels of Src detected with anti-pY416. Lower panels: Levels of total Src detected with anti-Src in same lysates. **(G,I)** Ratios of pY416 to total Src signals in lysates, normalized to control. (G) F_5,72_=4.464. DMSO vs. Brad., *P*=0.0167; DMSO vs. PMA, *P*=0.0226. (I) F_5,65_=11.06. DMSO vs. PMA, *P*=0.001; PMA vs. PP2+PMA, *P*<0.0001; DMSO vs. PP3+PMA, *P*=0.0042; PP3 vs. PP3+PMA, *P*=0.0086. **(J)** Upper panel: pY402 levels of Pyk2 detected with anti-pY402. Middle panel: pY579 levels of Pyk2 detected with anti-pY579. Lower panel: Levels of total Pyk2 detected with anti-Pyk2 in same lysates. **(K,L)** Ratios of pY402 and pY579 to total Pyk2 signals in lysates, normalized to control. (K) F_5,42_=35.85. DMSO vs. PMA, *P*<0.0001; PMA vs. PP2+PMA, *P*<0.0001; PP3 vs. PP3+PMA, *P*=0.0001; DMSO vs. PP3+PMA, *P*=0.0068; DMSO vs. PP2+PMA, *P*=0.0001; DMSO vs. PP2, *P*<0.0001. (L) F_5,68_=13.40. DMSO vs. PMA, *P*<0.0001; PMA vs. PP2+PMA, *P*<0.0001; PP3 vs. PP3+PMA, *P*=0.0362; DMSO vs. PP3+PMA, *P*=0.0202. **(B,C,E,G,I,K,L)** Data are presented as mean ± SEM. Number (*n*) of independent experiments for each condition are indicated inside bars. Statistical analysis was by ANOVA with post-hoc Bonferroni’s multiple comparisons test; \**P*≤0.05, \*\**P*≤0.01, \*\*\**P*≤0.001, \*\*\*\**P*≤0.0001. Bradykinin and PMA induced phosphorylation of Pyk2 on Y402 and Y579 and of Src on Y416, all of which were blocked by PF-431396 and PP2 but not the inactive PP3.

Importantly, PMA and bradykinin induced tyrosine phosphorylation of α_1_1.2 (Fig. 6). This effect was blocked by the Pyk2-inhibitor PF-431396 (Fig. 6A,B) and the Src inhibitors SU6656 and PP2, whereas the inactive PP2 analogue PP3 was without effect (Fig. 6C,D). These data show that activation of PKC translates into tyrosine phosphorylation of α_1_1.2 and that this requires both Pyk2 and Src.

**Figure 6.**
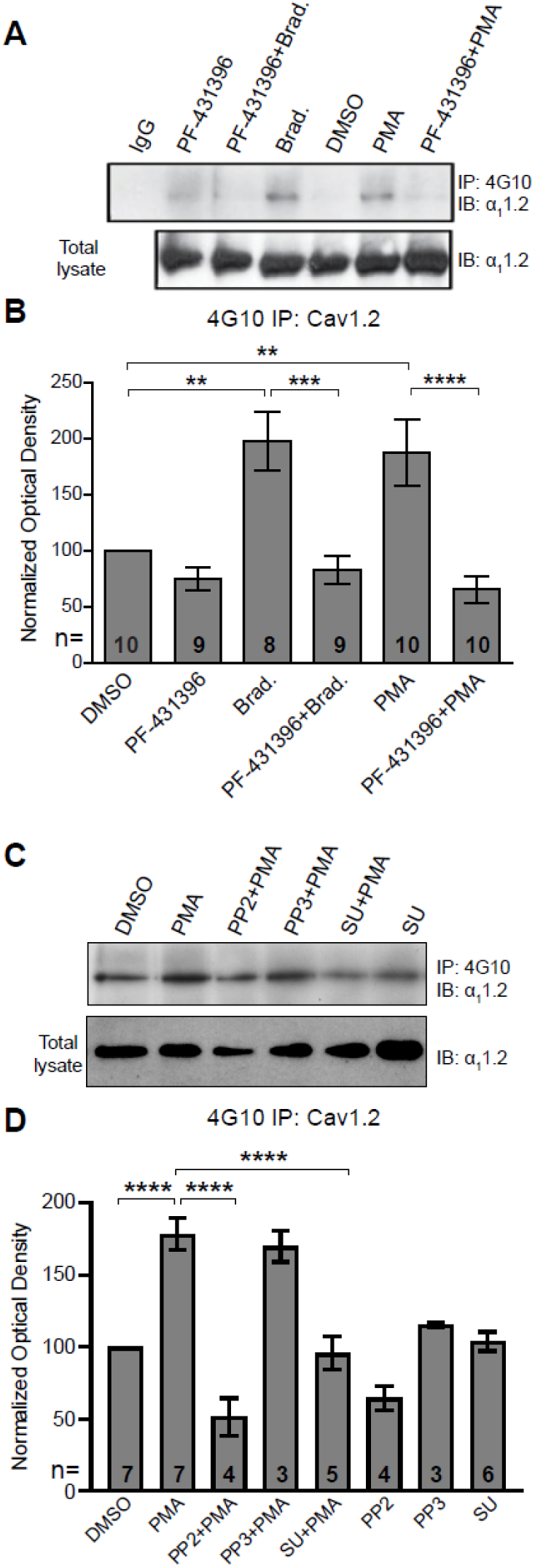
Increase in α_1_1.2 tyrosine phosphorylation by PKC is blocked by inhibitors or Pyk2 and Src. PC12 cells were treated as in Fig. 5 for analysis of tyrosine phosphorylation by IP with 4G10 and IB with anti-α_1_1.2. IgG indicates control IP with non-immune mouse IgG. The SU6656 was applied 5 min before PMA when indicated (SU, 10 μM). **(A,C)** Upper panels: pY of α_1_1.2 determined by 4G10 IP and α_1_1.2 IB. Lower panels: levels of total α_1_1.2 detected with anti-α_1_1.2 in corresponding lysates. **(B,D)** Ratios of pY signals in 4G10 IPs by IB with anti-α_1_1.2 to α_1_1.2 signals in lysates, normalized to control. Data are presented as mean ± SEM. Number (*n*) of independent experiments for each condition are indicated inside bars. Statistical analysis was by ANOVA with post-hoc Bonferroni’s multiple comparisons test. (B) F_5,50_=10.65. DMSO vs. Brad., *P*=0.0021; DMSO vs. PMA, *P*=0.0036; Brad. vs. PF-431396+Brad., *P*=0.0003; PMA vs. PF- 431396+PMA, *P*<0.0001. (D) F_7,31_=23.67. DMSO vs. PMA, *P*<0.0001; PMA vs. PP2+PMA, *P*<0.0001; PMA vs. SU+PMA, *P*<0.0001. (\*\**P*≤0.01, \*\*\**P*≤0.001, \*\*\*\**P*≤0.0001). Bradykinin- and PMA-induced α_1_1.2 tyrosine phosphorylation was blocked by PF-431396, SU6656 and PP2 but not the inactive PP3.

### Knock down of Pyk2 and Src prevents the increase in α_1_1.2 tyrosine phosphorylation upon stimulation of PKC

To control for any potential side effects of PF-431396 and determine whether Pyk2 is required for PKC-induced tyrosine phosphorylation of α_1_1.2 we employed FIV and HIV lentiviral expression vectors for shRNAs targeting Pyk2 in PC12 cells. We first designed and cloned an shRNA targeting rat Pyk2 (Sh1) into the FIV-based plasmid pVETL-GFP (*58, 84, 85*) and tested the ability and specificity of this construct to knockdown ectopically expressed Pyk2 in HEK293T/17 cells. Cells were co-transfected with vectors for expression of GFP-tagged rat Pyk2 (rPyk2-GFP) and the pVETL-Sh1-GFP or no shRNA control pVETL-GFP (Fig. 7A) and Pyk2 expression levels in the transfected cell lysates were examined via IB. Expression of rPyk2-GFP was virtually abolished by pVETL-Sh1-GFP whereas the control pVETL-GFP had no effect (Fig. 7A). IB with both tubulin and GAPDH antibodies confirmed that total protein levels were not affected by transfection of these plasmids (Fig. 7A).

**Figure 7.**
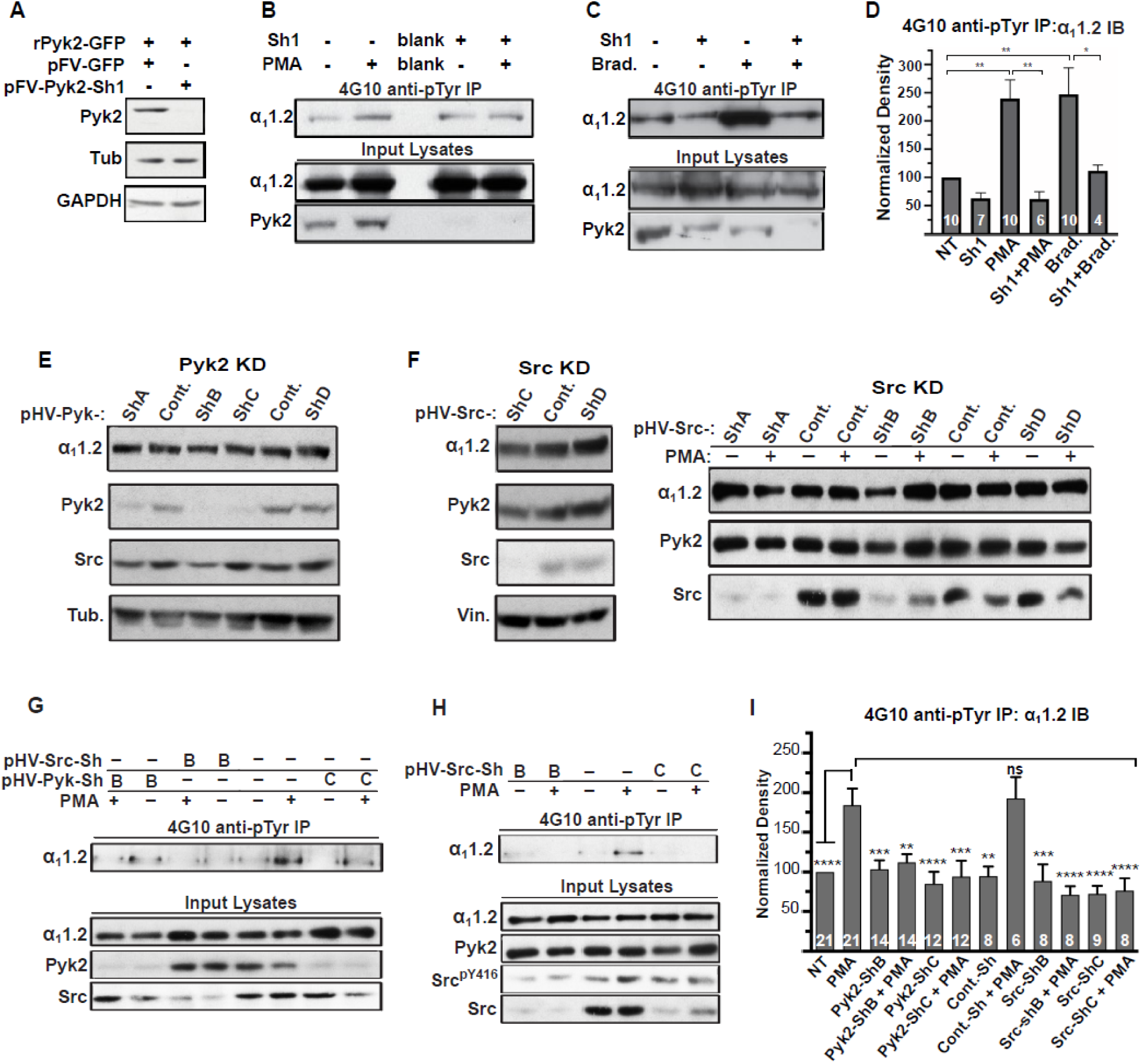
Increase in α_1_1.2 tyrosine phosphorylation by PKC is blocked by knockdown of Pyk2 and Src. **(A)** Lysates from HEK293T/17 cells transfected with vectors encoding rat Pyk2 (rPyk2-GFP) and either the Pyk2-targeting FIV lentivirus-derived, pVETL-Sh1-GFP (pFV-Pyk2-Sh1) or control (empty) pVETL-GFP (pFV-GFP) expression vectors, were immunoblotted (IB) with indicated antibodies. **(B,C)** IB analysis of indicated proteins in PC12 cultures incubated with viral particles containing pFV-Sh1-GFP (Sh1) FIV-based expression vector used in A or medium vehicle alone for 72h prior to treatment with either PMA (B), bradykinin (Brad.; C) or vehicle alone (-; B,C). Upper blots in B&C show anti-α_1_1.2 IBs of 4G10-anti-phosphotyrosine (pY) IP while middle and lower blots show direct IBs of indicated protein levels in input lysates. **(D)** Statistical analysis of the relative pY α_1_1.2 levels. F_5,41_=8.276. NT vs. PMA, *P*=0.0031; NT vs. Brad., *P*=0.0017; PMA vs. Sh1+PMA, *P*=0.001; Brad vs. Sh1+Brad, *P*=0.0433. **(E,F)** Direct IB analysis of indicated proteins in lysates of PC12 cultures transduced with HIV vector-derived lentiviral particles (e.g., pGFP-Pyk2-ShB-Lenti) containing expression cassettes for GFP and either the Pyk2-targeting (denoted pHV-Pyk-ShB and -ShC), Src-targeting (denoted pHV-Src-ShC and -ShD) or scrambled hairpin control (Cont.) shRNAs. In some cases (right blot) cultures were treated with PMA (+) or vehicle alone (-) before harvesting for IB. **(G,H)** IB analysis of indicated proteins from PC12 cultures infected with lentiviral particles containing HIV-GFP expression vectors as in E&F prior to treatment with either PMA (+) or vehicle (-). Upper panels show anti -α_1_1.2 IBs of 4G10-anti-pY IP while lower blots show direct IBs of input lysates with indicated antibodies. **(I)** Statistical analysis of relative α_1_1.2 pY levels. F_11,129_=6.180. NT vs. PMA, *P*<0.0001; PMA vs. Pyk2-ShB, *P*=0.0005; PMA vs. Pyk2-ShB+PMA, *P*=0.0029; PMA vs. Pyk2-ShC, *P*<0.0001; PMA vs. Pyk2-ShC+PMA, *P*=0.0002; PMA vs. Cont.-Sh, *P*=0.0019; PMA vs. Cont.-Sh+PMA, *P*>0.9999; PMA vs. Src-ShB, *P*=0.0007; PMA vs. Src-ShB+PMA, *P*<0.0001; PMA vs. Src-ShC, *P*<0.0001; PMA vs. Src-ShC+PMA, *P<*0.0001. The bar graphs in (D) and (I) show ratios of quantified anti-α_1_1.2 IB signals in 4G10 IPs relative to α_1_1.2 IB signals in total lysates, normalized to not treated (NT) control. Comparisons are made between samples treated with PMA (or bradykinin in D) and each of the other indicated conditions. Data are presented as mean ± SEM. Number (*n*) of independent experiments for each condition are indicated inside bars. Statistical analysis by ANOVA with post-hoc Bonferroni’s multiple comparisons test (ns= not significant vs. PMA*, *P*≤0.05, \*\**P*≤0.01, \*\*\**P*≤0.001, \*\*\*\**P*≤0.0001).

Next we tested whether pVETL-Sh1-GFP would inhibit PMA- and bradykinin-induced α_1_1.2 phosphorylation in PC12 cells. pVETL-Sh1-GFP lentiviral particles carrying the Sh1-shRNA and GFP expression cassettes were used to efficiently infect PC12 cells. Infected cells were monitored for GFP expression and then subjected to overnight serum starvation (to ensure low signaling levels) before treatment with PMA or bradykinin. Fully SDS-dissociated tyrosine- phosphorylated proteins were immunoprecipitated with 4G10 before SDS-PAGE and α_1_1.2 IB (Fig. 7B,C). As before, PMA and bradykinin induced a 2 to 2.5 fold increase in tyrosine phosphorylation of the α_1_1.2 subunit, which was strongly repressed by Sh1 (Fig. 7B-D). IB for total Pyk2 content confirmed Pyk2 knockdown by ∼70-90% (Fig. 7B,C bottom panels). These findings indicate that depletion of Pyk2 potently blunts the PKC-mediated increase in α_1_1.2 tyrosine phosphorylation. Total α_1_1.2 content was not altered by Sh1. These findings indicate that Pyk2 knockdown does not affect α_1_1.2 expression levels and that the reduction in tyrosine- phosphorylated α_1_1.2 is likely not due to any potential off-target effects of the pVETL-Sh1- shRNA.

To further verify and extend these findings we obtained HIV-GFP lentiviral vectors for expression of validated unique 29mer shRNAs targeting Pyk2 (HIV-GFP-Pyk2ShA-D) and Src (HIV-GFP-SrcShA-D) as well as a scrambled, non-silencing control (HIV-GFP-Shscr). HIV-GFP- Pyk2ShB and C and HIV-GFP-SrcShB and C were most effective in knocking down endogenous Pyk2 and Src, respectively (Fig. 7E,F and data not shown). PC12 cells were transduced with HIV-GFP-Pyk2ShB and C and HIV-GFP-Shscr, serum starved, stimulated with PMA, harvested, and lysed before 4G10 IP and IB for α_1_1.2, Pyk2, Src, and tubulin. Total protein levels of α_1_1.2, Pyk2, Src, and tubulin in lysate were monitored in parallel. The Pyk2- targeting HIV-GFP-Pyk2ShB and C but not the scrambled control shRNA abrogated the PMA- induced increase in α_1_1.2 tyrosine phosphorylation (Fig. 7G). Similarly, the Src targeting HIV- GFP-SrcShB and C but not the scrambled control shRNA blocked the increase in α_1_1.2 tyrosine phosphorylation upon PMA application (Fig. 7G,H). For quantification, phosphotyrosine signals were normalized to total α_1_1.2 in lysate (Fig. 7I). None of the HIV viral constructs exhibited any detectable effects on protein expression of α_1_1.2, Pyk2, Src or α-tubulin, vinculin, and GAPDH as determined in lysates suggesting these constructs did not affect general protein expression. Collectively, the above findings indicate that knockdown effects were specific and not simply the consequence of viral infection or expression of non-specific stem-loop RNAs. Basal levels of α_1_1.2 tyrosine phosphorylation were not significantly altered by sole application of any lentiviral particle preparation (Fig. 7D,I). Taken together, our findings strongly support the hypothesis that PKC signaling mediates its effects on Ca_V_1.2 through Pyk2 and Src.

### Inhibition of Pyk2 and Src blocks LTCC-dependent LTP

Ca_V_1.2 is concentrated in dendritic spines (*69, 86, 87*) where it mediates Ca^2+^ influx (*88, 89*) and several forms of LTP (*11, 12, 28, 29, 90*). Notably, in older mice and rats, about half of the LTP (called LTP_LTCC_) induced by four 200 Hz tetani, each 0.5 s long and 5 s apart, is insensitive to NMDAR blockage but abrogated by inhibition or elimination of Ca_V_1.2 (*28, 29, 91-93*).

Pharmacological inhibition and genetic disruption of Ca_V_1.2 also abolishes LTP induced by either pairing presynaptic stimulation with backpropagating action potentials (*90, 94, 95*) or by 5 Hz / 3 min tetani, the latter form of LTP requiring β_2_AR signaling to upregulate Ca_V_1.2 activity (*11, 12, 96*). Thus, we hypothesized that upregulation of Ca_V_1.2 activity by α_1_AR signaling can augment LTCC-dependent forms of LTP.

LTP_LTCC_ is prominent in mice older than 1 year (30-40% above baseline) but small in mice younger than 3 months (10-15% above baseline) (*91, 92*). LTP_LTCC_ requires Ca_V_1.2 activity (*29*) and stimulation of PKC signaling via type I metabotropic glutamate receptors (mGluR) (*93*). We tested whether increasing Ca_V_1.2 activity through α_1_AR - PKC - Pyk2 - Src - signaling can augment LTP_LTCC_. Similar to previous reports (*91, 92*), LTP_LTCC_ was ∼10% and was not statistically significant above baseline in our 13-20 week old mice (Fig. 8A). However, when Ca_V_1.2 activity was upregulated by stimulation of α_1_ARs with PHE, robust LTP_LTCC_ occurred (*P*≤0.05, Fig 8A). This LTP_LTCC_ was completely blocked by the LTCC inhibitor nimodipine and the α_1_AR antagonist prazosin (both *P*≤0.001, Fig 8A). Thus, this elevated potentiation strictly depends on both the activity of LTCCs and signaling through α_1_ARs. Importantly, this LTP_LTCC_ is also blocked by the Pyk2 inhibitor PF-719 and the Src inhibitor PP2 (both *P*≤0.001, Fig 8B).

**Figure 8.**
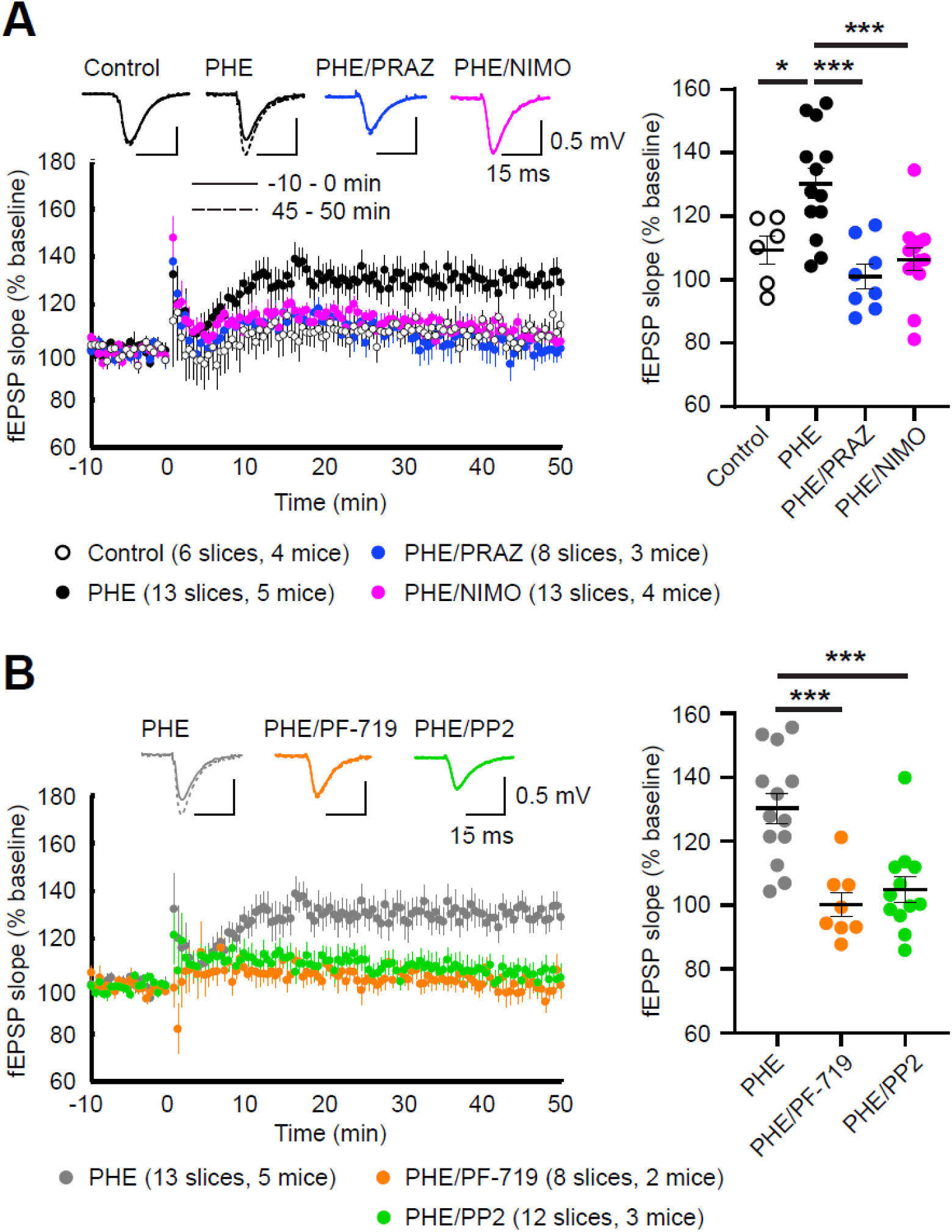
α_1_AR signaling augments LTP_LTCC_ through LTCC activity, Pyk2, and Src. LTP_LTCC_ was induced by four 200 Hz tetani, each 0.5 s long, in the CA1 Schaffer collateral projections to CA1 in acute hippocampal slices from 13-20 weeks old mice. **(A)** LTP_LTCC_ required PHE (10 μM) and was prevented by the LTCC blocker nimodipine (10 μM; NIMO) and the α_1_AR antagonist prazosin (1 μM; PRAZ). F_3,36_=9.937. Control vs. PHE, *P*=0.012; PHE vs. PHE/PRAZ, *P*=0.0001; PHE vs. PHE/NIMO, *P*=0.0003. **(B)** PHE-mediated LTP is blocked by inhibitors of Pyk2 (1 μM PF-719) and Src (10 μM PP2). F_2,30_=13.90. PHE vs. PHE/PF-719, *P*=0.0002; PHE vs. PHE/PP2, *P*=0.0003. Dot plots on the right show potentiation of fEPSPs determined as the averages of all responses between 45 and 50 min after high frequency stimulation (HFS) as % of averages of all responses in the 5 min preceding HFS. Bars and whiskers represent means ± SEM (\**P*≤0.05, \*\*\**P*≤0.001; one-way ANOVA with the Bonferroni correction). The number of slices and mice used are indicated.

These data indicate that robust LTP_LTCC_ in 13-20 week old mice requires Pyk2 and Src activity downstream of engaging α_1_AR to boost LTCC activation to sufficient levels.

## DISCUSSION

NE is arguably the most important neuromodulator for alertness and attention, augmenting multiple behavioral and cognitive functions (*1–5*). The G_q_ - coupled α_1_AR has the highest affinity among the cognate NE receptors and has been implicated in many studies in attention and vigilance (*16–22*). Inspired by our earlier findings that Ca_v_1.2 forms a unique signaling complex with β_2_AR, G_s_, AC, and PKA, making it a prominent effector of NE (*7, 11, 12*), we tested and found that Ca_v_1.2 is also a main target for NE signaling via the α_1_AR. Given that Ca_v_1.2 fulfills numerous functions in many cells this is a key and critical finding (*23, 24*). In the following paragraphs we discuss the four central and notable outcomes of our study.

Firstly, we found that the α_1_AR-selective agonist PHE strongly increased LTCC activity in neurons (Fig. 1). This increase was blocked by the α_1_AR-selective antagonist prazosin confirming that the PHE effect was mediated by the α_1_AR. Stimulation of two other major classes of G_q_-coupled receptors in neurons, mGluR1/5 and muscarinic M1/3/5 receptors, did not affect LTCC activity at the cell soma (Fig. S1). Accordingly, the α_1_AR augments LTCC activity through stimulation of a specific G_q_ – mediated signaling pathway rather than through general activation of G_q_. Thus, activity of these channels is precisely and judiciously regulated by α_1_AR signaling. More generally, these observations indicate that certain G_q_ signals can selectively target specific effector proteins. Defining what restricts the receptor type that can regulate to LTCCs will be an interesting avenue of future investigation.

Secondly, we identified a complex PKC/Pyk2/Src cascade that mediates regulation of Ca_v_1.2 by the α_1_AR. Clear evidence for this signaling pathway is provided by the inhibition of PHE-induced upregulation of LTCC activity by inhibitors of PKC, Pyk2, and Src (Fig. 2), which is further supported by the finding that direct stimulation of PKC also upregulates LTCC activity via Pyk2 and Src (Fig. 3). The role of the PKC/Pyk2/Src pathway in regulating Ca_v_1.2 is also substantiated by the association of Pyk2 in addition to Src and PKC with Ca_v_1.2 (Fig. 4) and inhibition of PKC-induced tyrosine phosphorylation of Ca_v_1.2 by Pyk2 and Src inhibitors (Fig. 6) and Pyk2 and Src knockdown (Fig. 7). Multiple shRNAs specifically targeting both Pyk2 and Src efficiently prevented the PKC-mediated increase in α_1_1.2 tyrosine phosphorylation. These observations not only confirm that Pyk2 mediates the Ca_v_1.2 regulation downstream of PKC but also indicates that Src itself is in this context the relevant member of the Src kinase family.

Thirdly, we found that Pyk2 is firmly associated with Ca_v_1.2 under basal conditions as reflected by their co-IP (Fig. 4). This association places Pyk2 into a complex that also contains its immediate upstream activator and downstream effector, i.e., PKC and Src. PKC can directly bind to the distal C-terminal region of α_1_1.2, which also contain S1928 (*66*). Given that S1928 is a phosphorylation site for PKC (*66*), it is conceivable that PKC binding to this region reflects a temporary kinase - substrate interaction rather than a more permanent association of PKC with Ca_v_1.2, although an interaction with a phosphorylation site does not rule out that PKC can stably bind to another region in the C-terminus of α_1_1.2. In addition, the A kinase anchor protein AKAP150, which is a major interaction partner for Ca_v_1.2 (*10, 79, 97*), binds not only PKA but also PKC (*98*) and constitutes another potentially constitutive link between PKC and Ca_v_1.2 (*67*). Furthermore, like Pyk2, Src co-precipitates with Ca_v_1.2 (Fig. 4) and binds directly to α_1_1.2 (*60, 65, 68*). Our determination that Pyk2 co-precipitates with Ca_v_1.2 from not only brain but also heart indicates that Ca_v_1.2 forms a signaling complex with Pyk2 and likely PKC and Src in various tissues.

We identified the loop between domains II and III of α_1_1.2 as the binding site for Pyk2. This observation lends further support to the association of Pyk2 with Ca_v_1.2. Future functional studies will test the relevance of this interaction for the regulation of Ca_v_1.2 by PKC and Src, either by acute application or expression of a peptide derived from this loop segment or mutational determination and abrogation of the peptide motifs required for Pyk2 binding. Of note, Src binds to residues 1955-1973 in rat brain α_1_1.2 (corresponding to residues 1982-2000 in the original rabbit cardiac α_1_1.2 (*99*)) (*65, 68*). This interaction with α_1_1.2 might bring Src in close proximity to loop II/III-associated Pyk2 to augment their structural and functional interaction once Pyk2 has been activated by PKC.

Formation of supramolecular signaling complexes or ‘signalosomes’ consisting of kinases and their ‘customers’ ensures fast, efficient, and specific signaling (*43, 70*). Our work establishes the PKC-Pyk2-Src-Ca_v_1.2 complex as such a signalosome. Furthermore, it defines how various G_q_- coupled receptors stimulate the activity of Ca_v_1.2 in different cells. The remarkably strong upregulation of Ca_v_1.2 channel activity by Src (*62, 65*) and upon activation of the PKC-Pyk2-Src signaling cascade as shown here rivals the upregulation by β adrenergic signaling, which is a central and thus widely studied mechanism of regulating Ca^2+^ influx into cardiomyocytes during the fight or flight response (*13, 100–106*). In fact, Ca_v_1.2 also assembles all components required for β adrenergic signaling including the β_2_AR, G_s_, adenylyl cyclase, and PKA in brain (*7, 79*) and heart (*13*). Formation of this complex is important for upregulation of Ca_v_1.2 activity (*11, 13*) and long-term potentiation of glutamatergic synapses induced by a 5 Hz theta rhythm during β adrenergic stimulation (*11, 12*). Analogously, assembly of the PKC-Pyk2-Src-Ca_v_1.2 may be important for fast and specific regulation of Ca_v_1.2 by the α_1_AR. This hypothesis can now be tested by pursuing determination of the precise binding site of Pyk2 in the loop between domains II and III of α_1_1.2 and then disrupting this interaction with peptides and point mutations. In rat brain neurons LTCC activity is increased by Src via phosphorylation of α_1_1.2 on Y2122 (*62, 65*). However, this phosphorylation site is not conserved even within rodents. It is equivalent to position 2150 in rabbit cardiac α_1_1.2, which is a Cys not Tyr residue (*99*).

Accordingly, other Tyr residues must serve as phosphorylation sites. It will be an interesting challenge for future work to identify the exact phosphorylation site and then test its functional relevance.

Remarkably, the Src inhibitor PP2 also completely blocked PKC-induced autophosphorylation of Pyk2 on Y402 (Fig. 5J,K). This finding indicates a close interdependence between Pyk2 and Src activation by PKC in PC12 cells. It is consistent with earlier results indicating that Pyk2 activation (assessed by Y402 phosphorylation) requires catalytically active Src (*107–110*), although in other systems Y402 phosphorylation was not dependent on Src (*81, 111, 112*).

Accordingly, Pyk2 autophosphorylation on Y402 and the consequent binding of Src to phosphoY402 induces Src-mediated phosphorylation of Pyk2 on Y579 or Y580 in its activation loop, which further enhances Pyk2 activity beyond the level achieved by Pyk2 autophosphorylation on Y402 (*55, 81, 113, 114*). Such interdependence was supported by the observation that Y579 phosphorylation upon PKC stimulation by either PMA or bradykinin was also completely blocked by the Src inhibitor PP2 (Fig. 5J, L).

Fourthly and finally, LTP_LTCC_ induced by 200 Hz tetani in 13-20 week old mice required stimulation of α_1_ARs and is completely blocked by inhibitors of LTCC, Pyk2, and Src. Incidentally, we were not able to induce any LTP_LTCC_ in mice younger than 13 weeks. Taken together, our findings suggest that LTP_LTCC_ requires stimulation of Ca_v_1.2 activity by α_1_AR - PKC - Pyk2 - Src signaling. While we focus here on the importance of α_1_AR signaling for LTP_LTCC_, this is not the only form of LTP that requires upregulation of Ca^2+^ influx through Ca_v_1.2.

Prolonged theta tetanus LTP (PTT-LTP), which is induced by a 3-minute-long 5 Hz tetanus, also depends on upregulation of Ca_v_1.2 activity (*28, 29, 92, 93, 115*). In PTT-LTP this upregulation is accomplished by β_2_AR - G_s_ - adenylyl cyclase/cAMP - PKA signaling and the ensuing phosphorylation of the central pore-forming α_1_1.2 subunit of Ca_v_1.2 on S1928 by PKA (*11, 12, 96*). Whether signaling by NE through α_1_AR and β_2_AR can act in parallel and is additive will be an interesting question for future studies. However, we already know that at least for classic PTT-LTP β_2_AR signaling is sufficient and does not require engagement of α_1_AR signaling (*96*). Because regulation of Ca_v_1.2 by β_2_AR signaling is highly localized (*7, 11*), it is conceivable that α_1_AR signaling might engage a subpopulation of Ca_v_1.2 channels whose spatial distribution differs from that of β_2_AR-stimulated Ca_v_1.2 in dendrites. Alternatively, parallel engagement of α_1_AR and β_2_AR signaling might ensure more robust and possibly additive or synergistic responses both at the Ca_v_1.2 channel level and in the synaptic potentiation that results.

LTP is thought to underlie learning and memory (*116, 117*). Conditional knock out of Ca_V_1.2 in the hippocampus and forebrain impaired LTP_LTCC_ as well as initial learning (*29*) and long-term memory of spatial Morris water maze tasks (*118*). Moreover, decreased Ca_V_1.2 expression or infusion of LTCC blockers into the hippocampus impaired both, LTP induced by pairing backpropagating action potentials in dendrites with synaptic stimulation and latent inhibition (LI) of contextual fear conditioning, the latter requiring learning to ignore non-relevant environmental stimuli (*90*). These Ca_V_1.2-related learning deficits might be in part due to impaired attention the animals pay to their experimental environment during learning phases, processes requiring concerted attention (*119*). Attention, in turn, depends on the neurotransmitter NE, which may well augment spatial learning through regulation of Ca_V_1.2 via α_1_AR - PKC - Pyk2 - Src signaling.

Ca_V_1.2 is increasingly implicated in not just the postsynaptic physiological functions discussed above. Multiple genome-wide association studies point to variants in the Ca_V_1.2 gene, *CACNA1C*, as major risk factors for schizophrenia, bipolar disorder, and other mental diseases (*23, 120–124*). Other studies link chronic upregulation of Ca_V_1.2 activity to etiologies behind senility and Alzheimer’s disease (e.g., (*125–128*)). Thus, it appears likely that dysfunctional regulation of Ca_V_1.2 contributes to these diseases, making the detailed molecular analyses of the signaling paradigms that regulate its functionality especially important to advance our mechanistic understanding for development of future therapies.

Here we establish for the first time that NE upregulates Ca**_V_**1.2 activity via a complex α_1_AR signaling cascade through PKC, Pyk2, and Src, the activity of each component being essential for LTP_LTCC_ and thus most likely relevant for learning. Our work forms the foundation for future studies to uncover the physiological context in which this action of NE is specifically engaged, what the precise role for each kinase is in this signaling cascade regulating Ca**_V_**1.2 activity, and how the individual kinases could be coordinately regulated to further fine-tune Ca**_V_**1.2 function. Given the central role of NE in attention and the many physiological and pathological aspects of Ca**_V_**1.2, regulation of this channel via NE - α_1_AR signaling predictably will elicit widespread and profound functional effects.

## MATERIALS AND METHODS

### MATERIALS AVAILABILITY STATEMENT

Further information and requests for resources and reagents should be directed to and will be fulfilled by the corresponding author, Johannes W. Hell.

### EXPERIMENTAL MODEL AND SUBJECT DETAILS

#### Animals

Pregnant Sprague-Dawley (SD) rats were ordered from Envigo (Order code 002) or Charles River (Strain code 001) and E18 embryos were used for preparation of dissociated hippocampal neuronal cultures. SD rats used for preparation of tissue extracts from heart and brain were of either sex and around 3 months old. For LTP experiments, mice of the strain B6129SF1/J aged between 13-18 weeks (both males and females) were used.

Animals were maintained with a 12-h/12-h light/dark cycle and were allowed to access food and water *ad libitum*. All procedures followed NIH guidelines and had been approved by the Institutional Animal Care and Use Committee (IACUC) at UC Davis (Protocol # 20673 and 22403).

#### Primary hippocampal neuronal cultures

Primary hippocampal neurons were maintained at 37°C in humidified incubators under 5% CO_2_ and 95% air. Both male and female rat embryos were used to prepare the cultures. Neurons were maintained in a medium containing 1x B-27 supplement (Gibco Cat#17504044), 1x Glutamax (Gibco Cat#35050061), 5% fetal bovine serum (FBS, Corning Cat#35-010-CV), and 1 µg/ml gentamicin (Gibco Cat#15710-064) in Neurobasal medium (Gibco Cat#21103-049). 10 µM each of 5-fluoro-2’-deoxyuridine (Sigma-Aldrich Cat#F0503) and uridine (Sigma-Aldrich Cat#U3003) were added around DIV7 to block the growth of glial cells.

#### Cell line

All cells were grown at 37°C in humidified incubators under 5% CO_2_ and 95% air. Rat pheochromocytoma cell line PC12 (ATCC Cat# CRL-1721; RRID:CVCL_0481, male) was grown in RPMI 1640 media (Gibco Cat#11875-101) containing 10% horse serum (HS, Gemini Bio Products Cat#100-508) and 5% FBS. For serum starvation, PC12 were incubated for 18 h in RPMI 1640 media containing 1% HS and 0.5% FBS. HT-1080 cells (ATCC CCL-121; RRID:CVCL_0317, male) used for virus titration were grown and maintained in MEM (Gibco Cat#11095-080) supplemented with 10% FBS. HEK293T/17 (ATCC Cat# CRL-11268, RRID:CVCL_1926, female) cells were routinely cultured in DMEM supplemented with 10% FBS (Gibco Cat#11995-065). Cell lines used were obtained from ATCC, expanded and frozen at low passage number. Care was taken during the use of cell lines to ensure that only one cell line was processed in the culture hood at a time, and that they were used within 25 to 30 passages. Their morphology in culture and doubling time were routinely monitored as were other distinguishing properties such as high transfectability (HEK293T/17 cells) or high Cav1.2 expression (PC12 cells) before the time of experimental use.

### METHODS DETAILS

#### Culture of primary hippocampal neurons

Hippocampal neurons were cultured from wild-type E18 male and female embryos from Sprague Dawley rats. Hippocampus was excised from the brains of embryos in ice-cold Hank’s Buffer (Sigma-Aldrich, Cat#H2387) with 10 mM HEPES (Gibco Cat#15630-080), 0.35 g/L NaHCO_3_ and 5 ug/ml gentamicin (Gibco Cat#15710-064) and digested in 0.78 mg/ml papain (Roche, Cat#10108014001) in 5 ml of the same buffer at 37°C for 30 min, in an incubator containing 5% CO_2_ and 95% air. Digested hippocampal tissue was washed with neuron medium twice, and triturated in the medium. The medium used for washes, trituration and culture of neurons consists of 1x B-27 supplement, 1x Glutamax, 5% FBS and 1 µg/ml gentamicin in Neurobasal medium (as stated in the section “Experimental Model and Subject Details”). 15,000 neurons were plated per well in 24-well plates on coverslips coated with poly-DL-ornithine (Sigma-Aldrich Cat#P0671) and laminin (Corning Cat#354232) and cultured in an incubator at 37°C and 5% CO_2_ and 95% air.

#### Single-channel recording

Single-channel recording was performed on hippocampal neurons on DIV10-15 using the cell-attached configuration at an Olympus IX50 inverted microscope as before (*11, 12, 129*). The membrane potential was fixed at ∼ 0 mV using a high K^+^ external solution. The external (bath) solution contained (in mM) 145 KCl, 10 NaCl, 10 HEPES and 30 D-glucose (pH 7.4 with NaOH, 325-330 mOsM). The internal (pipette) solution contained 110 mM BaCl_2_, 20 mM TEA-Cl, 10 mM HEPES, 500 nM BayK 8644 (Tocris Cat#1546; 200 nM used for Fig. 1-2; 500 nM used for Fig. 3) and 1 µM each of ω-conotoxins MVIIC and GVIA (China Peptides, custom synthesized) (pH 7.2 with TEA-OH, 325-330 mOsM). 3.5-5.5 MΩ resistance pipettes were used. The concentrations of kinase inhibitors, α_1_AR receptor blocker and agonist are as indicated in the figure legends. Neurons were preincubated with kinase inhibitors or receptor blocker in culture medium for 10 min prior to the experiment, and kinase inhibitors and receptor blocker were, where relevant, present in the external solution during recording. In experiments where isradipine was used, it was placed in the pipette solution, and no pre-incubation with isradipine was performed. Recordings started within 10 min of placing coverslips in the recording chamber in a bath solution with or without PHE (Sigma-Aldrich Cat#P6126), PMA (Merck Millipore Cat#524400), and different kinase inhibitors and receptor blocker. Currents were sampled at 100 kHz and low-pass filtered at 2 kHz using an Axopatch 200B amplifier (Axon Instruments) and digitized using Digidata 1440A digitizer (Axon Instruments). Step depolarizations of 2-s duration (one sweep) were elicited to the patch from – 80 mV to 0 mV at a start-to-start stimulation interval of 7 s. Typically, 100 sweeps were recorded per neuron and only cells with more than 70 sweeps recorded were analyzed. The single-channel search event detection algorithm of Clampfit 10.7.0.3 (Axon Instruments) was used to analyze open probabilities (NPo). Ensemble average traces was computed by averaging all sweeps from one neuron and averaging the averaged traces from all neurons in each group.

#### Co-immunoprecipitation of Cav1.2 with Pyk2 and Src

Brains and hearts were homogenized with a Potter tissue homogenizer in 10mL of a homogenization buffer containing 50 mM Tris-HCl (pH 7.4), 150 mM NaCl, 5 mM EGTA pH 7.4, 10 mM EDTA, 1% Triton-X 100, 25 mM NaF, 25 mM sodium pyrophosphate, 1 mM 4-nitrophenyl phosphate, 2 μM microcystin and protease inhibitors (1 μg/mL leupeptin (Millipore Cat#108975), 2 μg/mL aprotinin (Millipore Cat#616370), 10 μg/mL pepstatin A (Millipore Cat#516481) and 200 nM phenylmethylsulfonyl fluoride/PMSF). High-speed centrifugation was performed at 40,000 rpm for 30 min at 4°C. 500 μg of total brain or heart lysate extracts were incubated with 15 μL of Protein-A Sepharose beads (CaptivA protein resin, Repligen, Cat#CA- PRI-0100) and 2 μg of anti-Cav1.2 α1-subunit or control rabbit IgG antibody. Samples were incubated at 4°C for 4 h before being washed three times with ice-cold wash buffer (0.1% Triton X-100 in 150 mM NaCl, 10 mM EDTA, 10 mM EGTA, 10 mM Tris, pH 7.4). Samples were then resolved by SDS-PAGE and transferred onto PVDF membranes before immunoblotting with anti-Cav1.2 α1-subunit (J.W. Hell lab), -Pyk2 (Millipore Cat#05-488; RRID:AB_2174219), and - Src (J.S. Brugge lab) antibodies. The antibody dilutions used are listed in Table S2.

#### GST pulldown assay

Fragments of intracellular loops of Cav1.2 α_1_-subunit (Table S1) (*130, 131*) were expressed in *E. coli* strain BL21 as GST fusion proteins, purified, and integrity verified by IB essentially as previously described (*69, 132–134*). Overnight cultures from single colonies of the corresponding plasmids were cultured initially in 50 ml of LB medium containing ampicillin (100 μg/ml) with aeration until saturation. Incubation temperature was optimized for each expression construct to optimize translation and stability and varied between 28 and 37°C. After a 1:10 dilution into the same medium, cultures were grown for about 2-4 h until an A600 of about 0.8 was reached when isopropyl-β-D-thiogalactopyranoside was added for induction. After 4-5 h bacteria were collected by centrifugation (5000 rpm, SLA 3000 rotor, ThermoFisher Cat#07149) for 15 minutes and resuspended by gentle trituration in ice-cold 50 ml of TBS Buffer (150 mM NaCl, 15 mM Tris-Cl, pH 7.4) containing protease inhibitors (1 μg/ml pepstatin A, 1μg/ml leupeptin, 1μg/ml aprotinin, and 200 nM PMSF. 0.1 mg/ml lysozyme was added to lyse cell walls. The mixture was kept on ice for 30 min before addition of Sarcosyl (1.5% final concentration), β-mercaptoethanol (10 mM), and DNAse (50 U) for 15 minutes to fully solubilize the fusion proteins. In order to neutralize Sarcosyl, Triton X-100 was then added to a final concentration of 5%. Insoluble material was removed by ultracentrifugation (1h, 4°C, 40,000 rpm, Ti70 rotor, Beckman Coulter Cat#337922). The fragments were immobilized onto glutathione Sepharose (Millipore/Cytiva Cat#17-5132-02) for 3 h, washed three times with Buffer A (0.1% TX-100, 10 mM Tris-HCl, pH. 7.4) and incubated with affinity-purified His-tagged Pyk2 separately expressed in *E. coli* (3 h, 4°C). Beads were washed three times in Buffer A and bound proteins were eluted and denatured in SDS sample buffer, resolved by SDS-PAGE and transferred to a nitrocellulose membrane. IB with anti-Pyk2 antibody (Millipore Cat#05-488; RRID:AB_2174219) was used to detect Pyk2 binding during pull-down.

#### Analysis of Phosphorylation in PC12 cells

Drugs were used at the following concentrations: 2 µM PMA (Merck Millipore Cat#524400), 1 µM Bradykinin (Sigma-Aldrich Cat#05-23-0500), 3 µM PF-431396 (Tocris Cat#4278), 10 µM PP2 (Sigma-Aldrich Cat#P0042), 10 µM PP3 (Tocris Cat#2794), and 10 µM SU6656 (Sigma-Aldrich Cat#S9692).

For phospho-tyrosine analysis PC12 cells were washed after drug treatment twice in ice-cold PBS containing 1 mM pervanadate and 25 mM NaF. Samples were collected in PBS containing pervanadate, NaF, and protease inhibitors (see above), sonicated and extracted with SDS dissociation buffer (50 mM Tris-HCl, 1% SDS) at 65°C for 10 min. The SDS was neutralized with a fivefold excess of buffer A containing phosphatase and protease inhibitors. 500 μg of total protein from PC12 cell extracts were incubated over night at 4°C with 2 μg of the phospho- tyrosine 4G10 (Sigma-Alrich/Upstate Cat# 05-321; RRID: AB_2891016) or mouse control antibody (Jackson Immunoresearch Cat#015-000-003) and 15 μL of Protein-G Sepharose (Millipore/Cytiva Cat#GE-17-0618-05), washed three times in ice-cold wash buffer (0.1% Triton X-100 in 150 mM NaCl, 10 mM EDTA, 10 mM EGTA, 10 mM Tris, pH 7.4), resolved by SDS-PAGE and transferred onto PVDF membranes for IB. The antibody dilutions used are listed in Table S2. For re-probing, blots were stripped in 62.5 mM Tris-Cl, 20 mM DTT and 2% SDS at 50°C for 30 min. Chemiluminescence immunosignals were quantified using ImageJ (*135*) by multiple film exposures of increasing length to ensure signals were in the linear range (*128, 134*). Variations in total amounts of α_1_1.2, Pyk2, and Src in the different PC12 cell lysates was monitored by direct IB of lysate aliquots. Lysate signals was used to correct α_1_1.2 signals after 4G10 IP for such variations by dividing the latter by the former.

#### Lentiviral Constructs for shRNA to Pyk2 and Src

A list of all shRNA target sequences is provided in Table S3. The shRNA sequence Sh1 against Pyk2 has been validated for Pyk2 knockdown (*136*). The Sh1 sequence was cloned in the reverse orientation into the MfeI site of the lentiviral transfer vector pVETL-eGFP (*58, 84, 85*) for expression of Pyk2 shRNA and GFP to visualize infection. All HIV plasmids (HIV-GFP- Pyk2shA-D; HIV-GFP-SrcshA-D) for knocking down rat Pyk2 or Src as well as the scrambled, non-silencing hairpin control (HIV-GFP-shscr) were obtained from Origene (Cat# TL710108 and #TL711639). All expression plasmids (listed in Table S4) were confirmed by DNA sequencing.

#### Production of Lentivirus for Pyk2 and Src Knockdown

HEK293T/17 (ATCC Cat# CRL-11268, RRID:CVCL_1926) cells were plated onto 10 cm dishes at 1.8x10^6^ cells per dish and maintained until confluency (60-90%). Cells were transiently transfected with viral expression constructs using the calcium phosphate precipitation method (Jordan et al., 1996; Senatore et al., 2011). For FIV virus production, cells were transfected in a 3:2:1 ratio of parental vector (pVETL, FIV 3.2): pCPRD-Env: pCI-VSVG for a total of 24 μg of DNA per plate. For production of HIV viral particles targeting Src and Pyk2, cells were transfected at a ratio of 5:2:2:2 of parental vector (e.g. pGFP-Pyk2-shC-Lenti): pCI-VSVG: pMDL g/p RRE: pRSV-REV) according to manufacturer’s guidelines (OriGene). Media was exchanged 16 h after transfection. Media containing the packaged recombinant virus was collected at 48 h and 72 h, filtered through 0.45 μm filters, and concentrated by centrifugation (7,400 g for 16 h at 4°C). The viral pellet was resuspended in ice-cold PBS, aliquoted and stored at -80°C. Before use for transduction of PC12 cells all viral particle solutions were titered in the HT-1080 cell line (ATCC Cat#CCL-121; RRID:CVCL 0317) by seeding 12-well plates at 5x10^4^ cells per well one day before infection. 1, 5 and 10 μl of solutions containing concentrated HIV or FIV particles was added to each well and expression of GFP was monitored for 72 h post-infection before titer was calculated.

#### Slice preparation and electrophysiology

After decapitation, brains were removed from 13-18-week-old mice. 400 μm-thick transverse slices were made using a vibratome in cold, oxygenated (95% O_2_ and 5% CO_2_) dissection buffer (in mM: 87 NaCl, 2.5 KCl, 1.25 NaH_2_PO_4_, 26.2 NaHCO_3_, 25 glucose, 0.5 CaCl_2_, 7 MgCl_2_, 50 sucrose). Slices were allowed to recover at room temperature for at least 1 hr in oxygenated ACSF (in mM: 119 NaCl, 3 KCl, 2.5 CaCl_2_, 1.25 NaH_2_PO_4_, 1.3 MgSO_4_, 26 NaHCO_3_, 11 glucose). Following recovery, slices were transferred to a recording chamber and maintained at 32-33°C in oxygenated ACSF. Field excitatory postsynaptic potential (fEPSP) was evoked by stimulating the Schaffer collateral pathway using bipolar electrode, and synaptic responses were recorded with ACSF-filled microelectrodes (1-10 MΩ) placed in the stratum radiatum of CA1 region. Recordings were acquired using an Axoclamp-2B amplifier (Axon Instruments) and a Digidata 1332A digitizer (Axon Instruments). Baseline responses were collected at 0.07 Hz with a stimulation intensity that yielded 40–50% of maximal response. LTP was induced by four episodes of 200 Hz stimulation (0.5 sec) with 5-sec-intervals. To measure LTCC-mediated LTP, 50 μM D-APV (Tocris Cat#0106) was included in ACSF. When used the inhibitors (10 μM nimodipine (Bayer Charge: BXR4H3P), 1 μM prazosin, 1 μM PF-719 (*137*), and 10 μM PP2) were added to ACSF from the start of the recording. Phenylephrine (PHE, 10 μM) was added after at least 15 min of stable baseline, and LTP was induced ∼10 min after the addition of PHE.

### QUANTIFICATION AND STATISTICAL ANALYSIS

Statistical analyses were performed using Prism 5 or 9 (GraphPad). Data are presented as mean ± SEM. Sample sizes, *P* values and statistical tests are indicated in the figure legends. Statistical significance was determined using ANOVA with post-hoc Bonferroni’s multiple comparisons test (Fig. 5-7). For analysis of channel open probability (NPo, Fig. 1-3, Fig. S1-2), first outliers were identified using iterative Grubb’s method inbuilt in Prism. Then, statistical significance was determined using one-way ANOVA with post-hoc Holm-Sidak’s multiple comparisons test. For analysis of LTP (Fig. 8) a one-way ANOVA was applied followed by Bonferroni correction.

## DATA AVAILABILITY STATEMENT

Requests for raw datasets should be directed to and will be fulfilled by the corresponding author, Johannes W. Hell.

## ACKNOWLEDGEMENTS

We thank Dr. Stephen Strittmatter (Yale University) for providing PF-719.

## Funding

This work was supported by: National Institutes of Health grant R01 MH097887 (JWH) National Institutes of Health grant R01 AG055357 (JWH) National Institutes of Health grant R01 HL098200 (MFN) National Insitutes of Health grant R01 HL121059 (MFN), and National Institutes of Health grant T32 GM099608 (PBH).

## Author contributions

Conceptualization – KNMM, MCH, JWH

Performed final experiments – KNMM, PBH, KK, MS, MZ

Performed pilot experiments – MNC, MFN

Data analysis and visualization – KNMM, PBH, KK, MS, MZ, MCH

Supervision – MCH, JWH

Writing – original draft: KNMM, MCH, JWH

Writing – review and editing: KNMM, MNC, MFN, MCH, JWH

## Competing interests

Authors declare that they have no competing interests.

## Supplementary Figures and Tables

**Fig. S1.**
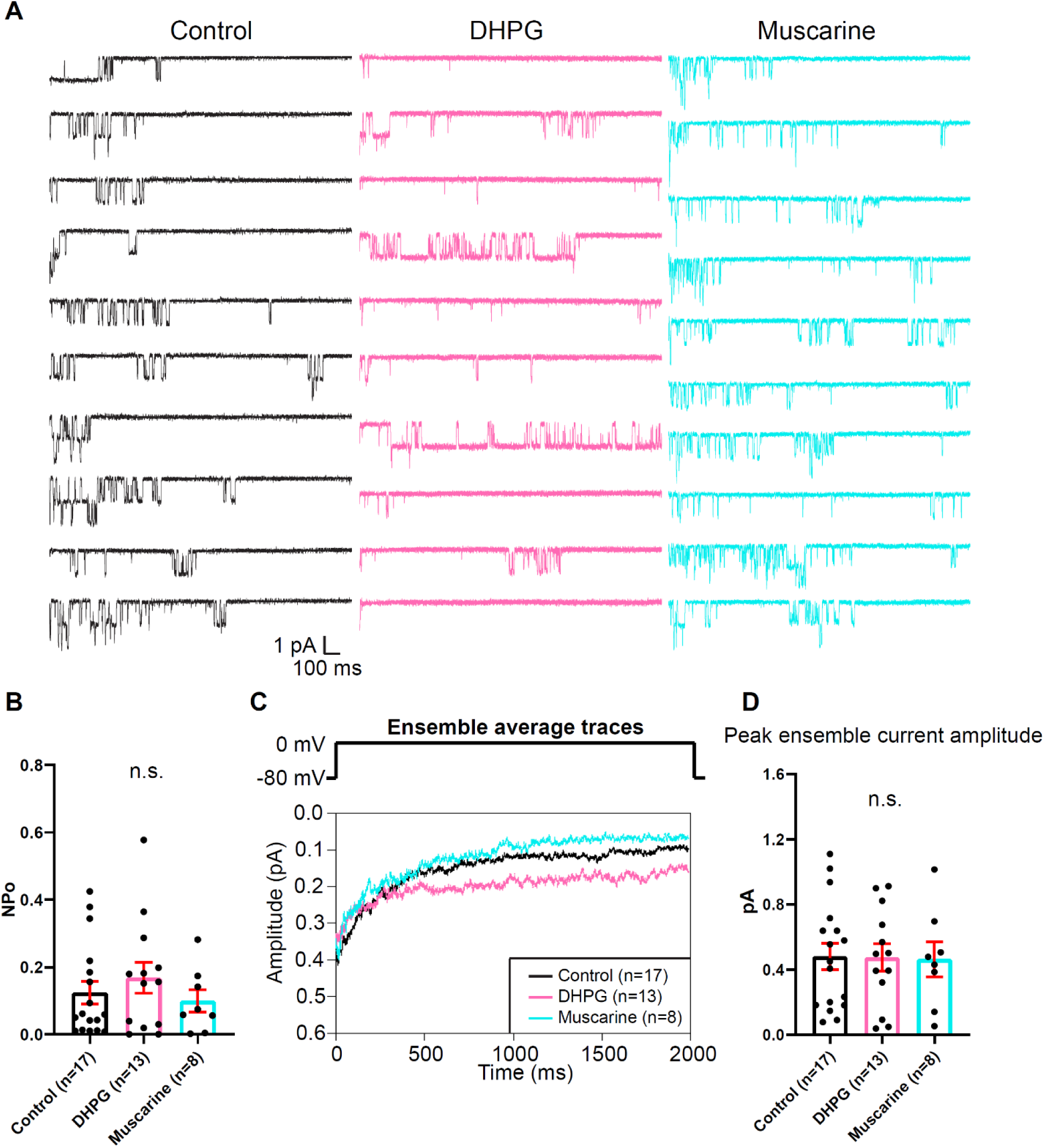
Group I mGluR and muscarinic receptor agonists did not change NPo of LTCCs in hippocampal neurons. **(A)** Ten consecutive traces from representative cell-attached single-channel recordings of LTCCs from cultured hippocampal neurons with vehicle (H_2_O; black), 100 µM DHPG (pink), 10 µM muscarine (cyan). **(B)** DHPG and muscarine did not alter NPo. F_2,35_=0.6559. Control vs. DHPG, *P*=0.648; Control vs. Muscarine, *P*=0.6843. **(C)** Ensemble averages during depolarization. **(D)** DHPG and muscarine did not alter the amplitudes of the peak ensemble average current. F_2,35_=0.007689. Control vs. DHPG, *P*=0.9904; Control vs Muscarine, *P*=0.9904. **(B,D)** Data are presented as means ± SEM. n represent the number of cells (n.s., not significant; ANOVA with post-hoc Holm-Sidak’s multiple comparisons test).

**Fig. S2.**
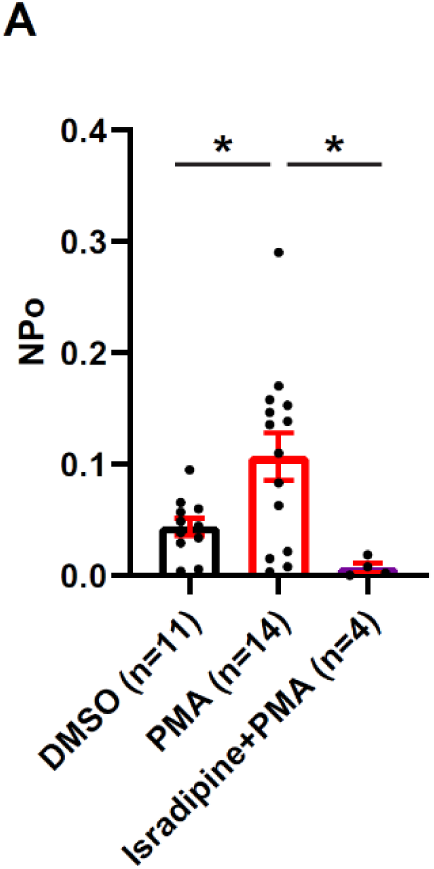
L-type channel blocker isradipine completely blocks L-type single-channel currents in the presence of PMA. PMA (2 μM) in the bath solution potentiated the open probability of L-type channels during single-channel recording. Isradipine (10 μM) in the pipette solution completely blocked L-type currents in PMA-containing bath solution. Data are presented as as means ± SEM. n represents the number of cells (\**P*≤0.05; ANOVA with post-hoc Holm-Sidak’s multiple comparisons test). F_2,26_=6.004. DMSO vs. PMA, *P*=0.0139; PMA vs. Isradipine+PMA, *P*=0.0129.

**Table S1.** Amino acid residues of fragments of intracellular loops of Cav1.2 α_1_-subunit used in GST pull-down studies.

| **Designation** | **Residues\*** | **Species** |
| --- | --- | --- |
| N-term | 2-124 | rat |
| Loop I-II | 409-526 | rat |
| Loop II-III | 754-901 | rat |
| Loop III-IV | 1165-1219 | rat |
| C-term | 1584-2140 | rat |
| CT-1 | 1507-1733 | rabbit |
| CT-23 | 1622-1905 | rabbit |
| CT-4 | 1909-2171 | rabbit |

\* residue number corresponds to the initial $\alpha_11.2$ sequence, which originated from rabbit heart (Mikami et al., 1989) (Gene Bank Accession number: CAA33546, NM\_001136522); rat numbers correspond to $\alpha_11.2$ as originally cloned from brain (Snutch et al., 1990) (Gene Bank Accession number: NM\_012517).

**Table S2.** Antibody dilutions used for immunoblotting.

| **Antibody** | **Used at dilution** |
| --- | --- |
| Anti-Cav1.2 α1-subunit (raised in the lab; RRID: AB_2891029) | 1:10000 |
| Anti-Pyk2 (Millipore Cat#05-488; RRID:AB_2174219) | 1:5000 |
| Anti-Pyk2 phospho-Y402 (Millipore Cat# 07-892;<br>RRID:AB_568885) | 1:20000 |
| Anti-Pyk2 phospho-Y579 (Life Technologies Cat#44-632G) | 1:10000 |
| Anti-Src (gift from J.S. Brugge) | 1:500 |
| Anti-Src phospho-Y416 (Millipore Cat# 05-857;<br>RRID:AB_441924) | 1:10000 |
| Anti-α-tubulin (Santa Cruz Biotechnology, Cat# sc32293; RRID:<br>AB_628412) | 1:40000 |
| Anti-GAPDH (Millipore/Cytiva, Cat# MAB374;<br>RRID:AB_2107445) | 1:10000 |

**Table S3.**
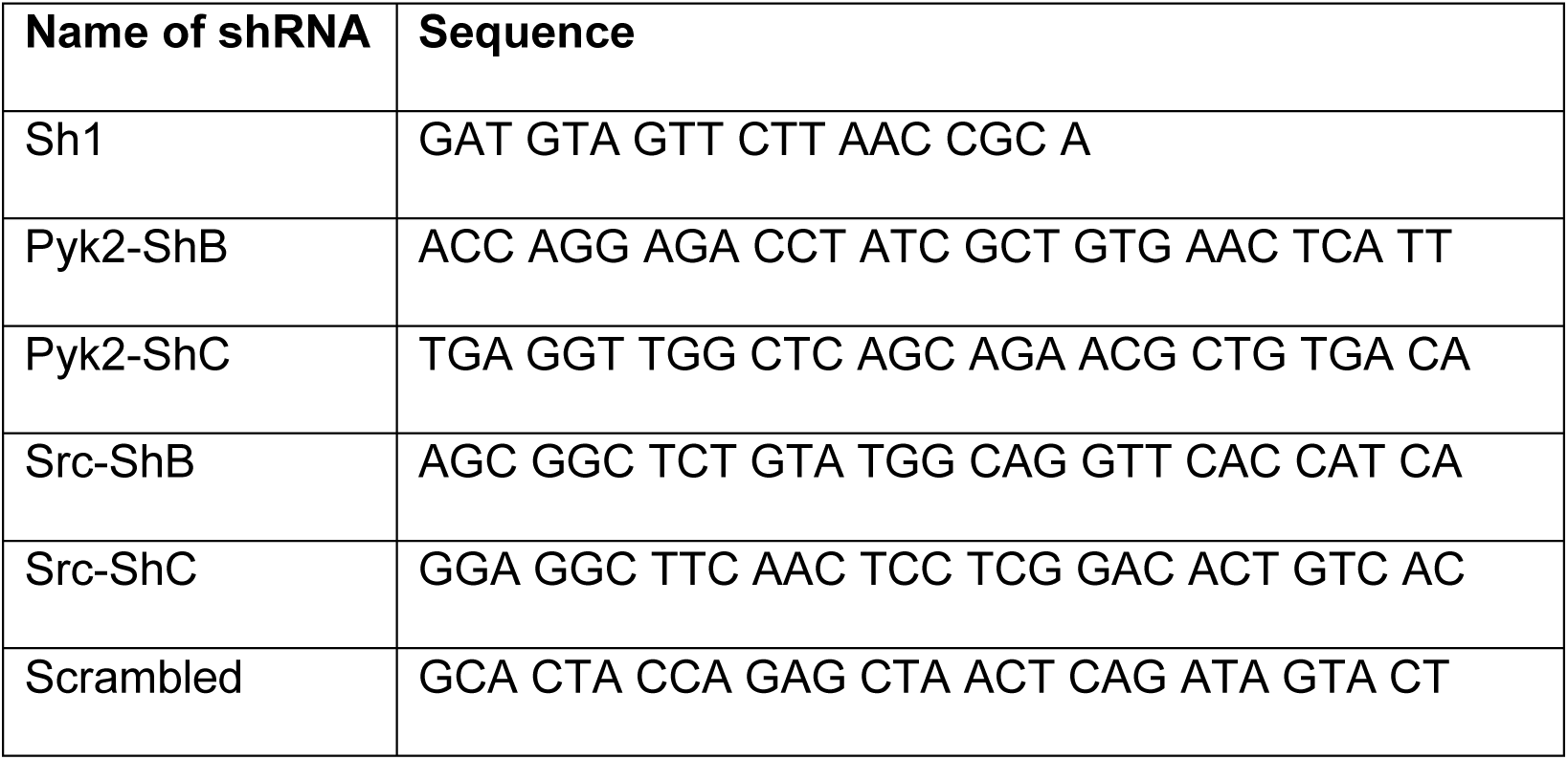
shRNA sequences targeting rat Pyk2 and Src.

**Table S4.**
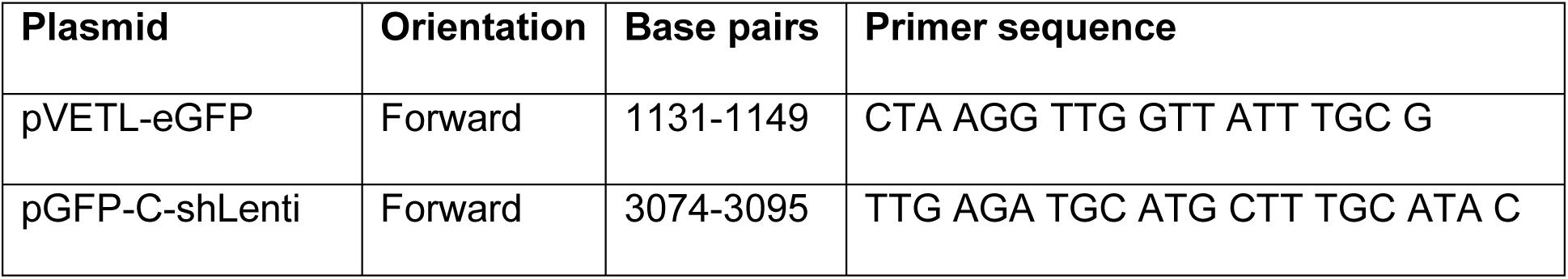
Sequencing primers for validation of knockdown constructs.

## Notes

### Competing Interest Statement

The authors have declared no competing interest.

